# Cellular taxonomy and spatial organization of the ventral posterior hypothalamus reveals neuroanatomical parcellation of the mammillary bodies

**DOI:** 10.1101/2020.05.14.096818

**Authors:** Laura E. Mickelsen, William F. Flynn, Kristen Springer, Lydia Wilson, Eric J. Beltrami, Mohan Bolisetty, Paul Robson, Alexander C. Jackson

**Affiliations:** Department of Physiology and Neurobiology, University of Connecticut, Storrs, CT, USA; Connecticut Institute for the Brain and Cognitive Sciences, Storrs, CT, USA; The Jackson Laboratory for Genomic Medicine, Farmington, CT, USA; Department of Genetics and Genome Sciences, University of Connecticut Health Center, Farmington, CT, USA; Institute for Systems Genomics, University of Connecticut, Farmington, CT, USA; National Institute of Diabetes and Digestive and Kidney Diseases (NIDDK), Bethesda, MD, USA; Bristol-Myers Squibb, Pennington, NJ, USA

**Keywords:** Hypothalamus, mammillary nuclei, tuberomammillary nucleus, mammillothalamic tract, anterior thalamus, single cell RNA-seq, cell types, neuropeptides, neurotransmitters

## Abstract

The ventral posterior hypothalamus (VPH) is an anatomically complex brain region implicated in arousal, reproduction, energy balance and memory processing. However, neuronal cell type diversity within the VPH is poorly understood, an impediment to deconstructing the roles of distinct VPH circuits in physiology and behavior. To address this question, we employed a droplet-based single cell RNA sequencing (scRNA-seq) approach to systematically classify molecularly distinct cell types in the mouse VPH. Analysis of >16,000 single cells revealed 20 neuronal and 18 non-neuronal cell populations, defined by suites of discriminatory markers. We validated differentially expressed genes in a selection of neuronal populations through fluorescence *in situ* hybridization (FISH). Focusing on the mammillary bodies (MB), we discovered transcriptionally-distinct clusters that exhibit a surprising degree of segregation within neuroanatomical subdivisions of the MB, while genetically-defined MB cell types project topographically to the anterior thalamus. This single cell transcriptomic atlas of cell types in the VPH provides a detailed resource for interrogating the circuit-level mechanisms underlying the diverse functions of VPH circuits in health and disease.

## INTRODUCTION

The ventral posterior hypothalamus (VPH) is a functionally and cytoarchitecturally complex region of the hypothalamus, dominated by the mammillary bodies (MB), a discrete diencephalic structure on the basal surface of the VPH. Surrounding VPH subregions include the caudal arcuate nucleus (Arc), caudal lateral hypothalamic area (LHA) and the tuberomammillary (TMN), premammillary (PM) and supramammillary (SUM) nuclei. These subregions are embedded within diverse neural systems, with widespread afferents and efferents, known to regulate diverse physiological and behavioral functions ^1–3^. The MB are best known as the diencephalic branch of the classic limbic circuit of Papez ^4^, linking the hippocampal formation with the anterior thalamus and midbrain, and are critical for spatial memory in both rodents and primate models, as well as episodic memory in humans ^5–9^. VPH subregions surrounding the MB are also robust modulators of behavioral state. The ventral and dorsal PM nuclei are implicated in reproductive and defensive behaviors respectively ^10–12^. The SUM is associated with arousal and modulation of theta rhythms ^13–15^. The Arc is a crucial node in the regulation of hunger, satiety and reproduction ^16–18^. Finally, the TMN, populated by histamine (HA)-synthesizing neurons, is an important modulator of wakefulness ^19–21^.

The functional diversity of the VPH is likely explained by cellular heterogeneity among these neuronal populations, the neural circuits they give rise to and the complex brain-wide networks in which they are embedded. A significant obstacle in understanding the circuit-level mechanisms underlying its function is that VPH neuronal cell type diversity is poorly understood. Here, we employed single cell RNA-sequencing (scRNA-seq) to develop a comprehensive molecular census of transcriptionally distinct cell types in the VPH of both male and female juvenile mice. In our unsupervised analysis of over 16,000 isolated single cells, we identify 18 non-neuronal clusters and 20 distinct neuronal clusters, the majority of which are glutamatergic. Transcriptionally distinct neuronal populations are defined by the unique expression of discriminatory markers that include neuropeptides, transcription factors, calcium-binding proteins and signaling molecules. We went on to validate differentially expressed genes in a selection of identified neuron populations through multiplexed fluorescent *in situ* hybridization (FISH), ISH data from the Allen Brain Institute (ABA) ^22^, as well as anterograde tract-tracing in genetically-distinct populations in the MB. Taken together, our identification of the population structure and cellular diversity of VPH cell populations provides a resource for detailed genetic dissection of VPH circuits and interrogation of their roles in behavior, in both health and disease.

## RESULTS

### Isolation of single cells from mouse VPH for transcriptomic analysis

In order to isolate single cells from the mouse VPH for scRNA-seq analysis, we microdissected the region of the VPH from fresh brain slices obtained from a total of five male and five female C57BL/6 mice (P30-34), similar to previously described procedures (Mickelsen et al, 2017; Mickelsen et al, 2019), in two independent harvests (see Materials and Methods). Single cell suspensions were loaded onto a 10X Genomics Chromium Controller (Fig. 1a) and processed using the 10X Genomics platform ^23^. All VPH microdissections were mapped for consistency using anatomical landmarks across the rostrocaudal axis (Fig. 1b) ^24^. The two harvests used 10X V2 and V3 chemistry, respectively, and both consisted of separate male and female pools for a total of four separate pools. We found little sex-dependent differences within each harvest (Fig. 1c) and samples from the two harvests were pooled with batch correction to account for chemistry-dependent differences (Fig. S1; see Materials and Methods). In our pooled data set, the median transcripts (UMIs)/cell was 8336 and the median genes/cell was 3415 (Fig. 1d). We used the 1,500 genes with highest normalized dispersion for dimensionality reduction using PCA and UMAP followed by cluster identification using Leiden community detection, identifying a total of 20 clusters in the first iteration of clustering (Figs. 1e and f). Neuronal and non-neuronal clusters were segregated using a two-component Gaussian mixture model trained on the per-cluster average expression of four pan-neuronal markers *Snap25, Syp, Tubb3, Elavl2* (Fig. S1b and c) leading to a binary classification of neuronal and non-neuronal cells (Figs. 1e and f). Subsequent clustering of only neuronal cells (20 clusters; Fig. S2a and b) and only non-neuronal cells (18 clusters; Fig. S2c and d) showed comparable proportions from each sex and batch.

**Figure 1:**
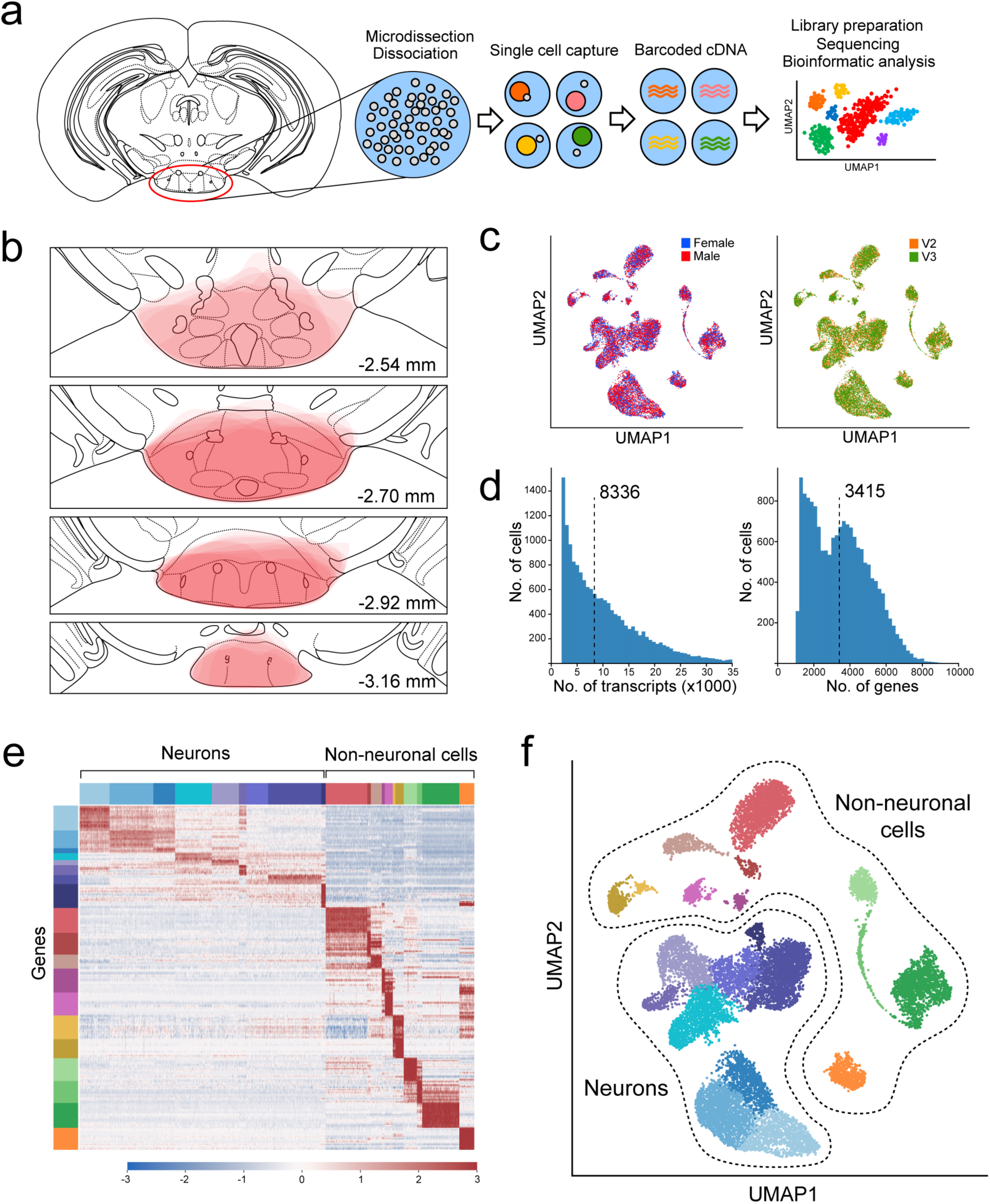
Overview of VPH microdissection, single cell isolation, batch correction and clustering. **(a)** Workflow schematic representing the VPH microdissection from coronal brain slices, single-cell dissociation, sequencing library preparation, and bioinformatic analysis. Adapted from ^44^. **(b)** Location of VPH microdissections mapped onto the coronal mouse brain atlas at distances from bregma of −2.54, −2.70, −2.92, −3.16 mm. Atlas images modified from Paxinos and Franklin (2012). **(c)** Two-dimensional UMAP plots representing 16,991 single cells from four sequencing libraries color coded by mouse sex (left) and 10X Genomics chemistry version (right) following batch correction. **(d)** Histograms of unique transcripts (left) and genes (right) detected in 16,991 single cells after quality control. Dashed vertical lines represent the median transcripts and genes per cell, respectively. **(e)** Heatmap and (f) UMAP plot showing the first iteration of unsupervised clustering revealing 20 unique clusters. Neuronal populations are disjoint from non-neuronal populations.

### Major non-neuronal cell types in the VPH

Among non-neuronal cell populations in the VPH, we identified a total of 18 distinct clusters distinguished from one another by suites of cell type-specific discriminatory markers (Fig. S3a-d). We resolved six distinct populations of oligodendrocyte lineage cells arranged in a contiguous strand in the UMAP plot (Fig. S3a), likely reflecting a developmental gradient of gene expression. These include oligodendrocyte precursor cells (OPC or NG2+ cells) (cluster 1), immature oligodendrocytes (cluster 2) and four oligodendrocyte populations (clusters 3-6), all of which exhibit a wave of differentially-expressed genes (ex. *Cspg4, Fyn, Ctps, Tspans2, Apod, Klk6*, etc.) (Fig. S4a and b) which aligned well with the diversity of functional markers of the oligodendrocyte lineage in the mouse brain identified through previous scRNA-seq analyses ^25–27^. We also resolved three distinct clusters of astrocytes (clusters 7, 8 and 9), all of which were *Aqp4*+ and *Agt*+. Clusters 7 and 8 were distinguished by high expression of *Slc7a10* and *Htra1* and low expression of *Gfap*, while cluster 9 exhibited high expression of both *Gfap* and *C4*b (Fig. S4c and d). In nearby clusters we identified tanycytes (cluster 10: *Rax*+) and ependymal cells (cluster 11: *Ccdc153*+, *S100a6*+) (Fig. S4c and d). In addition, we found other distinct clusters readily identifiable as macrophages (cluster 12: *Mrc1*+), microglia (cluster 13: *Tmem119*+), pericytes (cluster 14: *Rgs5*+), vascular smooth muscle cells (VSMCs) (cluster 15: *Rgs5*+, *Acta2*+), two populations of putative vascular leptomeningeal cells or VLMCs (cluster 16: *Fxyd5+, Slc47a1*+) (cluster 17: *Dcn*+) and vascular endothelial cells (VECs) (cluster 18: *Pecam1*+, *Slc38a5*+) (Fig. S3b). Our identification of major non-neuronal VPH cell types is based on, and is closely aligned with previous scRNA-seq analyses from mouse brain ^25–27^.

### Diverse populations of excitatory and inhibitory neuronal cell clusters in the VPH

Among the neuronal clusters, which contained significantly more genes and UMIs (4,334 and 12,065, respectively) per cell than the overall data set, we first examined broad patterns in the expression of fast amino acid and monoamine neurotransmitter markers (Fig. 2a). Expression of genes necessary for the synthesis/packaging of the excitatory transmitter glutamate (*Slc17a6*, encoding the vesicular glutamate transporter 2, VGLUT2) and the inhibitory transmitter GABA (*Slc32a1*, encoding the vesicular GABA transporter, VGAT) provided a binary classification of *Slc17a6*+ clusters as glutamatergic (VPH ^GLUT^) and *Slc32a1*+ clusters as GABAergic (VPH^GABA^) neurons. This was further supported by the expression of the gene encoding a synthetic enzyme for GABA (*Gad1*/GAD67) which largely aligned with *Slc32a1*+ clusters. We found that of the 20 neuronal clusters we identified, thirteen were glutamatergic, six were GABAergic and one (cluster 17) best matched the profile of a unique population of histaminergic (HA) neurons based on the unique expression of histidine decarboxylase (*Hdc*/HDC) (Fig. 2a and b). Overall, within these neuronal populations, clusters were distinguished by suites of differentially expressed transcripts (Fig. 2c, d) with comparable UMIs/cell and genes/cell (Fig. 3e). In the following analyses, we validated the co-expression of key markers and their spatial organization in selected VPH neuronal populations. In all cases, we first attempted to map transcriptionally distinct cell clusters onto specific anatomical subregions within the VPH, by comparing differentially-expressed transcripts with the online database of *in situ* hybridization (ISH) data from the Allen Brain Atlas (ABA) ^22^, followed by co-expression analysis using multiplexed FISH.

**Figure 2:**
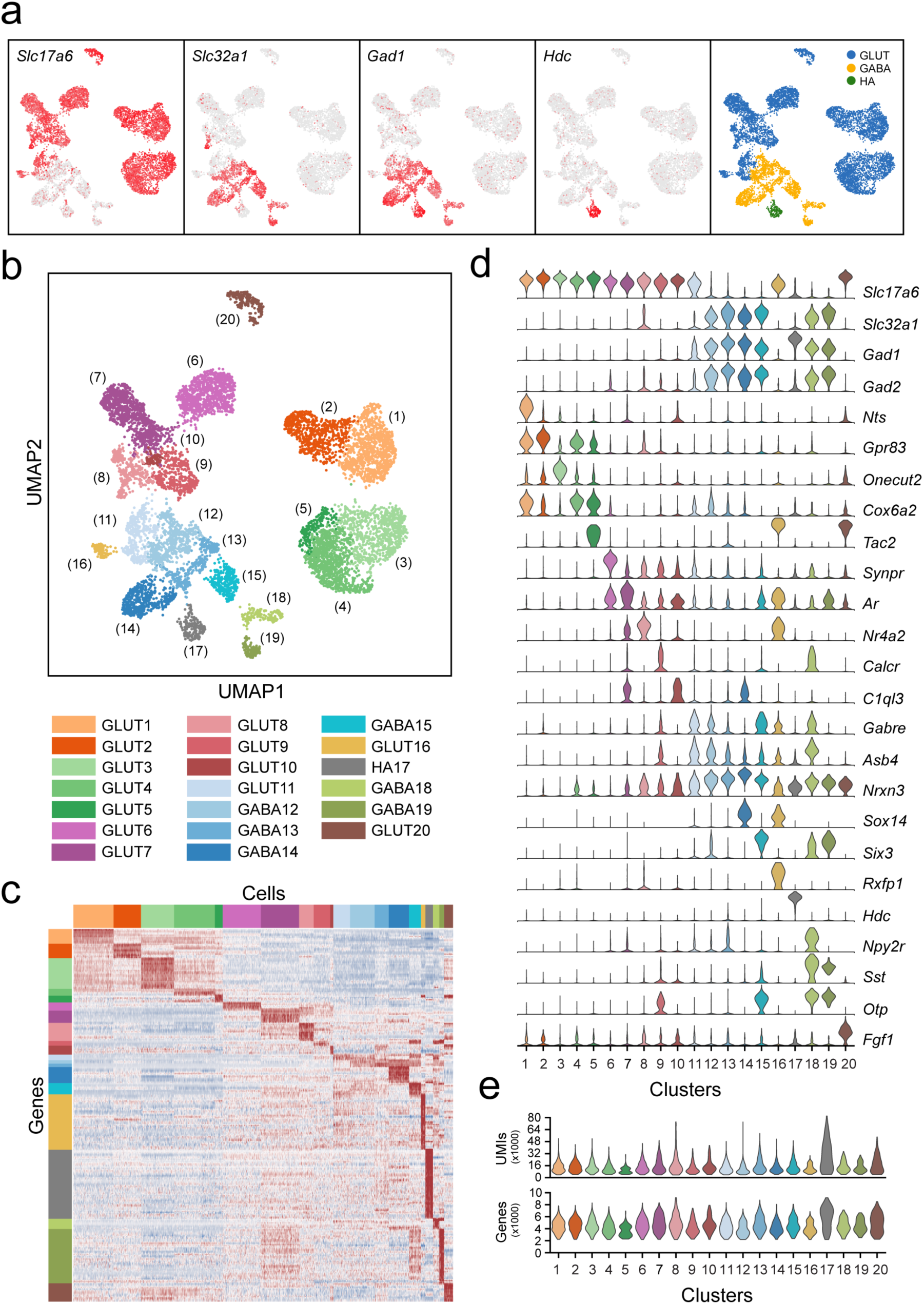
Classification of VPH neuronal populations. **(a)** UMAP plots showing normalized expression of *Slc17a6, Slc32a1, Gad1*, and *Hdc* after the second iteration of unsupervised clustering on just neuronal cells. Using these four genes, neurons were classified by a three-class Gaussian mixture model as glutamatergic (Glut, blue), GABAergic (GABA, yellow), or histaminergic (HA, green). **(b)** Unsupervised clustering of 20 VPH neuronal cell types shown in a UMAP embedding. **(c)** Heatmap showing scaled expression of discriminatory genes across all 20 neuronal clusters. **(d)** Violin plots showing the distribution of normalized expression in each cluster of neurotransmitters (*Slc17a6, Slc32a1, Gad1, Gad2*) (upper) and discriminatory marker genes (lower). **(e)** Violin plots showing the distribution in the number of unique transcripts (upper) and number of genes (lower) in each neuronal cluster.

**Figure 3:**
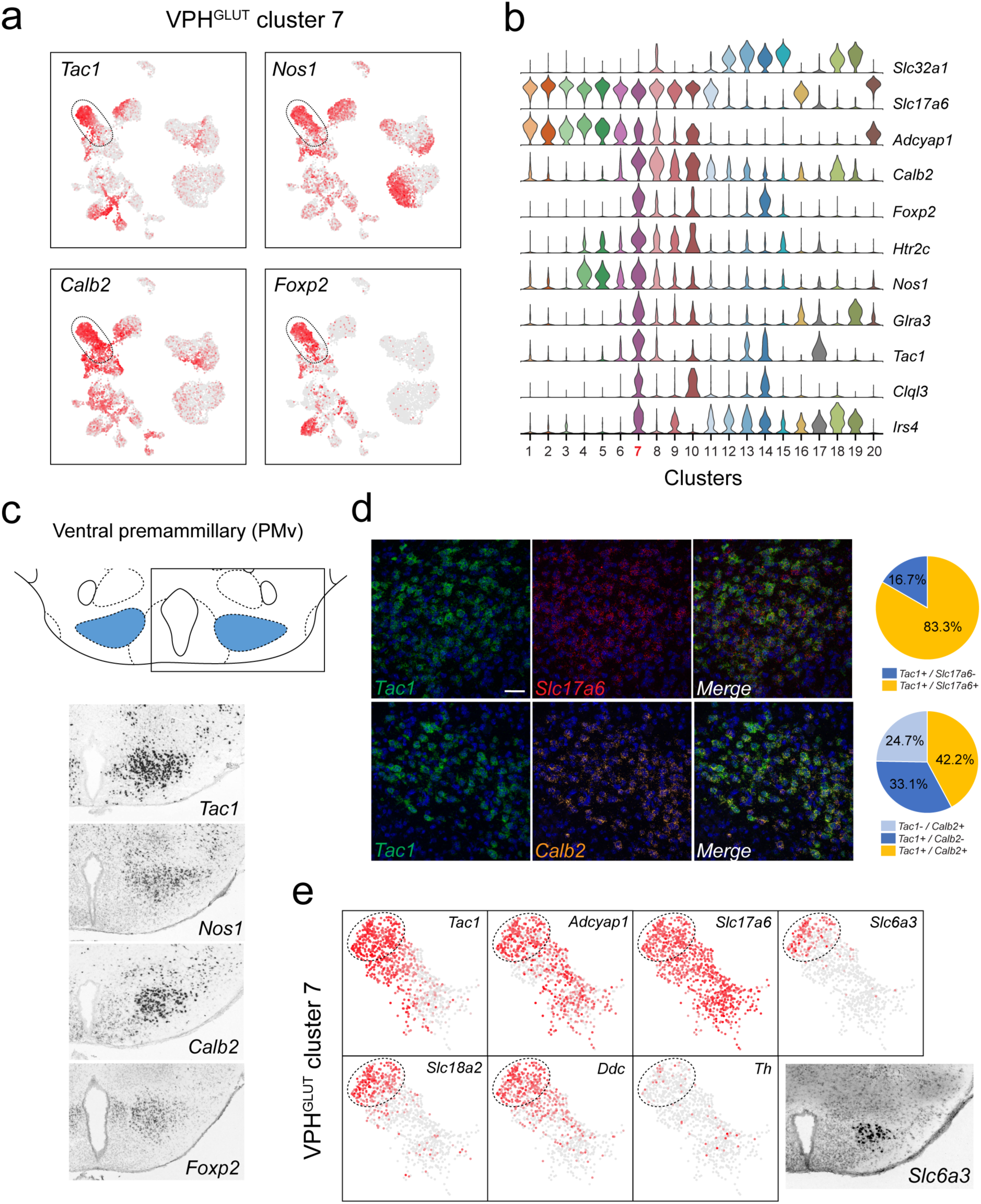
Identification of a population of putative *Tac1*+ PMv neurons and a *Slc6a3*+ PMv subpopulation. **(a)** UMAP plots showing normalized expression of *Tac1, Nos1, Calb2* and *Foxp2* enriched in VPH^GLUT^ cluster 7 following *Slc32a1* and *Slc17a6* (top). **(b)** Violin plot showing discriminatory marker genes enriched in cluster 7. **(c)** Mouse brain atlas schematic, modified from Paxinos and Franklin, 2012, showing the PMv in a coronal section at distance from bregma of −2.46 mm (top). ISH images for *Tac1, Nos1, Calb2* and *Foxp2* from the ABA ^22^ (bottom). **(d)** Confocal micrographs (40x) of coronal sections of wild type mice and corresponding pie charts representing co-expression of mRNA for *Tac1* and *Slc17a6* (n= 678 cells, 3 mice; upper) and *Tac1* and *Calb2* (n=963 cells, 3 mice; lower). Scale bar (applicable to all micrographs) 50 μm. **(e)** UMAP plots showing normalized expression of markers in VPH^GLUT^ cluster 7 only, including cell type markers for reference (*Tac1, Adcyap1* and *Slc17a6*) and markers that define a subpopulation of putative catecholaminergic neurons (*Slc6a3, Slc18a2, Ddc* and *Th*). ISH image from the ABA ^22^ showing *Slc6a3* expression in the PMv (inset).

### Transcriptional signatures differentiate ventral and dorsal premammillary nuclei

Among neuronal clusters that correspond to discrete anatomical subregions of the VPH, we identified at least one VPH^GLUT^ population with a suite of differentially-expressed transcripts that appears to closely match the ventral premammillary nucleus (PMv) (Fig. 3). VPH^GLUT^ cluster 7 expresses the following key transcripts: *Tac1* (encoding substance P or SP), *Nos1* (encoding neuronal nitric oxide synthase or NOS), *Calb2* (encoding calretinin) and *Foxp2* (encoding the transcription factor forkhead box P2) (Fig. 3a) along with a host of other differentially expressed markers (Fig. 3b). The spatial pattern of expression of each of the markers in Fig. 3a were found in the ABA (Lein et al, 2007), and correspond closely to the PMv (Fig. 3c). Through FISH co-expression analysis, we found that *Tac1, Calb2* and *Slc17a6* were extensively co-expressed in a cluster of neurons in the PMv (Fig. 3d). These data broadly align with several known markers for the PMv ^11^. For example, SP ^28,29^ and NOS ^30^ are both enriched in the PMv. Another marker that is enriched in VPH^GLUT^ cluster 7 is *Adcyap1* (encoding the neuropeptide pituitary adenylate cyclase-activating polypeptide, or PACAP) (Fig. 3b). Consistent with the role of the PMv in reproductive function ^11,12^, PACAP+ PMv neurons were recently shown to critically regulate female reproductive physiology and fertility ^31^.

Interestingly, we also found that a distinct subpopulation of VPH^GLUT^ cluster 7 neurons robustly expresses markers of catecholaminergic neurotransmission including *Slc6a3* (encoding the dopamine transporter, DAT), *Ddc* (encoding DOPA decarboxylase) and *Slc18a2* (encoding the vesicular monoamine transporter 2, VMAT2) but low expression of *Th* (encoding tyrosine hydroxylase) (Fig. 3e). This cluster corresponds well to a previously identified catcholaminergic ^32^, *Slc6a3*+ PMv population ^33^ that was more recently found, through circuit and behavioral analysis, to regulate male social behavior ^34^ and aggression ^35^ in a glutamate-dependent, but dopamine-independent, manner ^34^, consistent with the profile we identified (*Slc17a6*+, *Slc18a2*+, *Slc6a3*+, *Ddc*+, *Th*-). These transcriptomic data provide further biological insight into the repertoire of receptors and signaling molecules expressed by this key behavioral node.

Another distinct VPH^GLUT^ population (cluster 6) shared a number of common markers with clusters 1-5 (ex. *Cck, Foxb1*) that, in turn were largely undetectable in clusters 7-10 (Fig. S5a and b). Curiously, VPH^GLUT^ cluster 6 also expressed a suite of markers that were enriched in clusters 7-10 (ex. *Ebf3, Dlk1, Synpr, Nxph1*) that were largely undetectable in clusters 1-5 (Fig. S5a and b). In particular, *Synpr* (encoding the presynaptic protein synaptoporin), *Dlk1* (encoding delta like non-canonical Notch ligand 1), *Ebf3* (encoding early B cell factor 3) and *Stxbp6* (encoding the synaptic protein syntaxin binding protein 6) are enriched in VPH^GLUT^ cluster 6 (Fig. S5a). Examining the expression patterns of *Synpr, Dlk1* and *Stxbp6* in the ABA ^22^, we found that all three are enriched in a discrete region that appears to correspond well to the dorsal PM (PMd) and largely undetectable in the more caudal mammillary bodies (MB) (Fig. S5c, d). This suggests that VPH^GLUT^ cluster 6 likely represents a PMd population that expresses a number of unique signatures (ex. *Synpr, Stxbp6*) but shares some commonalities with VPH^GLUT^ clusters 1-5 (ex. *Foxb1, Cck, Adcyap1*) and VPH^GLUT^ cluster 7, a putative PMv population (ex. *Nxph1, Ar*).

### Transcriptional signatures that define a putative dual phenotype population in the supramammillary nuclei

Another notable neuronal cluster was VPH^GLUT^ cluster 8. Although all cells in cluster 8 express *Slc17a6*, and nominally classified as glutamatergic, a subpopulation appears to co-express *Slc32a1* and *Gad2* but not *Gad1* (Fig. S6a). Subjecting this cluster to another iteration of unsupervised clustering revealed six sub-clusters. One of these (GLUT8_3) closely corresponds to the *Slc17a6*+/*Slc32a1*+/*Gad2*+ subcluster and co-expresses a suite of discriminatory makers including *Sema3c, Inhba, Rxfp1* and others (Fig. S6b). Cross-referencing with the ABA ^22^ shows selective expression of *Sem3c, Inhba* and *Rxfp1* in the SUM (Fig. S6c, d). Interestingly, a unique population of VGLUT2/VGAT co-expressing axons originating in the SUM was found among projections to the hippocampal dentate gyrus ^36,37^ and these dual phenotype SUM neurons were found to indeed co-release GABA and glutamate onto neurons of the dentate gyrus ^38,39^. Our identification of this VPH^GLUT^ cluster 8 subpopulation is consistent with dual phenotype SUM neurons and provides further molecular insight into the biology of this unique population in addition to possible strategies for precise genetic targeting.

### Neuronal populations in the caudal arcuate and tuberal nuclei

A highly distinct neuronal cluster, VPH^GLUT^ cluster 16, expresses a suite of markers that identify it as a well-described neuronal cell type in the arcuate nucleus (Arc) - kisspeptin-neurokinin B-dynorphin (‘KNDy’) neurons (Fig. S7). KNDy neurons are considered to be the gonadotropin-releasing hormone pulse generator controlling release of luteinizing hormone from the anterior pituitary, and therefore essential for fertility and reproduction ^18,40,41^. We confirm that that this cluster co-expresses the defining neuropeptides *Tac2* (encoding neurokinin B), *Pdyn* (encoding dynorphin), *Kiss1* (kisspeptin), as well as the NKB receptor *Tacr3* (encoding the neurokinin 3 receptor) (Fig. S7a, b). The key markers that define KNDy neurons in our data set are consistent with those identified in previous mouse scRNA-seq data sets, both from whole hypothalamic samples ^42^ and Arc-specific samples ^43^. Consistent with the important role of Arc KNDy neurons in reproduction and fertility, we found high expression of the following hormone receptors: *Prlr* (encoding the prolactin receptor), *Esr1* (encoding the estrogen receptor 1), *Ar* (encoding the androgen receptor) and *Pgr* (encoding the progesterone receptor). Other markers that exhibit relatively robust and unique expression in KNDy neurons include *Nhlh2, Inhbb, Nr5a2, Rxfp1* and *Col2a1* (Fig. S7b). A selection of key markers for VPH^GLUT^ cluster 16 neurons were confirmed using ISH data from the ABA ^22^ (Fig. S7c) and correspond to the caudal Arc.

Another cluster, VPH^GABA^ cluster 18, also expresses a suite of markers that are known to be enriched in other Arc populations. While all neurons in VPH^GLUT^ cluster 18 express *Otp* and *Dlk1*, one subcluster likely corresponds to well-known Arc AGRP/NPY neurons (*Agrp* and *Npy*), and another likely corresponds to a previously identified population of Arc *Sst*+ neurons such as *Sst*/*Unc13c* ^43^. *Pomc* expression was more diffuse but concentrated in VPH^GLUT^ cluster 11 (Fig. S7d). Yet another cluster, VPH^GABA^ cluster 14 expresses a set of markers (*Dlk1, Thrb, Rxrg* and *Unc13c*) that appear to correspond well to a population of cells in the caudal Arc. In particular, using ISH data from the ABA ^22^, both *Dlk1* and *Thrb* (encoding the thyroid hormone receptor beta) are enriched in a cluster surrounding the 3^rd^ ventricle/mammillary recess corresponding to the medial posterior portion of the Arc ^24^ or posterior portion of the periventricular nucleus (Allen Institute Mouse Brain Reference Atlas).

Finally, two other distinct GABAergic neuronal populations appear to correspond to the tuberal region of the LHA, also known as hypothalamic tuberal nuclei. VPH^GABA^ cluster 15 (enriched in *Sst, Six3, Otp, Col25a1, Ecel1* and *Parm1*), and VPH^GABA^ cluster 19 (enriched in *Sst, Otp, Dlk1, Pthlh* and *Ptk2b*) (Fig. S7d), correspond closely with two previously identified subpopulations of *Sst*-expressing GABAergic neurons in the tuberal LHA (LHA^GABA^ clusters 6 and 13 respectively) ^44^. The most caudal region of the tuberal LHA would be expected to be included in the rostral-most sections of our VPH microdissections. *Sst*-expressing neurons are enriched in the tuberal region ^45^ and have recently been implicated in the regulation of feeding behavior in mice ^46^.

### Histaminergic neurons in the tuberomammillary nuclei (TMN)

A highly distinct cluster emerged from our dataset that exhibited a pattern of gene expression characteristic of histamine (HA)-producing neurons (VPH^HA^ cluster 17). HA neurons are a well-characterized monoaminergic, neuromodulatory population with cell bodies concentrated in the ventral TMN (TMNv) and dorsal TMN (TMNd), and also scattered throughout the VPH region. TMN HA neurons are the sole source of neuronally-synthesized HA in the brain, make widespread projections throughout the brain ^47,48^ and have a key role in modulating wakefulness ^19–21^. Previous mouse hypothalamic scRNA-seq studies have also detected a population of HA neurons ^42,43^. We found that VPH^HA^ cluster 17 neurons express the following key transcripts: *Hdc*, in addition to *Slc18a2* (encoding the vesicular monoamine transporter 2 or VMAT2), *Maob* (encoding monoamine oxidase B) and another unique marker Wif1 (encoding Wnt inhibitory factor 1) (Fig. 4a). Additional makers include transcripts for *Maoa, Msrb2, Itm2a, Bsx, Hrh3, Hcrtr2, Sncg* and *Prph* (Fig. 4b). Our identification of *Prph* (encoding the neurofilament protein peripherin) as a highly selective marker for HA neurons is consistent with previous anatomical work ^49^. Key markers (*Hdc, Slc18a2, Wif1* and *Maob*) correspond well to the TMNv and TMNd in the ABA ^22^ (Fig. 4c). FISH analysis of *Wif1* and *Maob* co-expression with *Hdc* revealed extensive co-expression in both the TMNv and TMNd (Fig. 4d). With regard to fast neurotransmitter phenotype, we found that VPH^HA^ cluster 17 neurons were exceptional in our dataset in that they express very low levels of both *Slc17a6* or *Slc32a1*, but robustly express *Gad1*. That TMN HA neurons are GAD+, and likely capable of GABA synthesis, has long been recognized ^50,51^. While there has been recent conflicting data on whether or not HA neurons express *Slc32a1* ^52,53^, our data indicate that TMN HA neurons robustly co-express *Gad1* accompanied by very low co-expression of *Slc32a1*.

**Figure 4:**
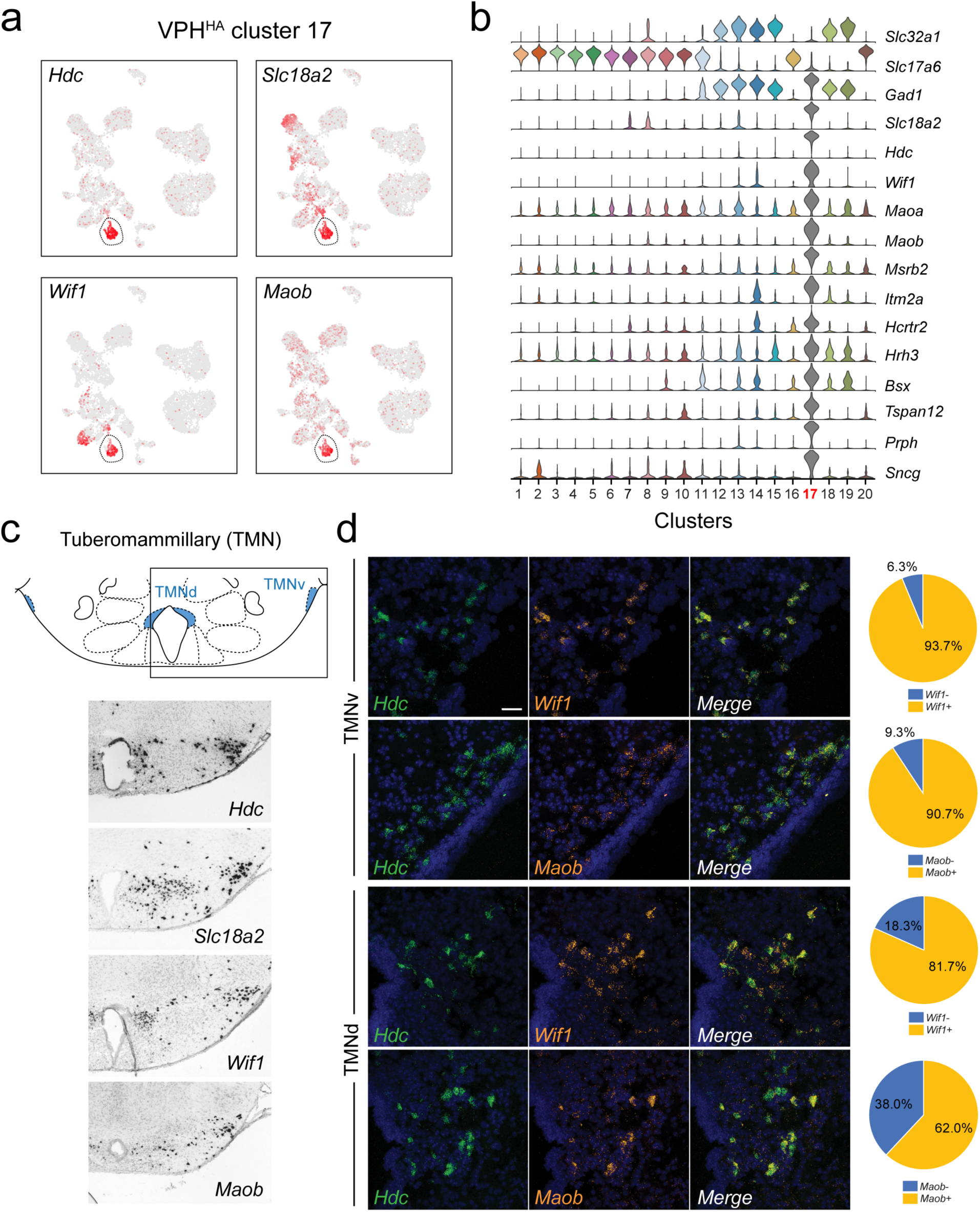
Identification of *Hdc*+ histaminergic (HA) neurons in the TMN. **(a)** UMAP plots showing normalized expression of *Hdc, Slc18a2, Wif1* and *Maob* enriched in VPH^HA^ cluster 17 (circled). **(b)** Violin plot showing discriminatory marker genes enriched in VPH^HA^ cluster 17 following *Slc32a1* and *Slc17a6* (top). **(c)** Mouse brain atlas schematic, modified from Paxinos and Franklin, 2012, showing the dorsal TMN (TMNd) and ventral TMN (TMNv) in a coronal section at distance from bregma of −2.54 mm (top). ISH images for *Hdc, Slc18a2, Wif1*and *Maob* from the ABA ^22^ (bottom). **(d)** Confocal micrographs (40x) of coronal sections of wild type mice and corresponding pie charts representing co-expression of mRNA in the TMNv (top) for *Hdc* and *Wif1* (n=415, 3 mice; upper) and *Hdc* and *Maob* (n=570, 3 mice; lower) and in the TMNd (bottom) for *Hdc* and *Wif1* (n=109, 3 mice; upper) and *Hdc* and *Maob* (n=192, 3 mice; lower). Scale bar (applicable to all micrographs) 50 μm.

### A distinct neuronal cluster corresponds to the lateral mammillary (LM) nuclei

We identified a distinct VPH^GLUT^ population (cluster 20) characterized by the expression of *Tac2, Pvalb* (encoding the calcium-binding protein parvalbumin), *Cplx1* (encoding the synaptic protein complexin-1) and *Tcf4* (encoding transcription factor 4) (Fig. 5a), in addition to a suite of other discriminatory markers including *Fgf1, Cbln2, Infg2, Nefm, Myo1a* and *Syt2* (Fig. 5b). The spatial pattern of expression of *Tac2, Pvalb, Cplx1* and *Tcf4* in the ABA ^22^ corresponds well to the lateral mammillary (LM) nuclei, which are tightly circumscribed bilateral cell clusters, immediately lateral to the medial mammillary region and bisected by the fornix (Fig. 5c). LM neurons are components of the MB and are implicated in the encoding of head direction and aspects of spatial ^5,8,54^. VPH^GLUT^ cluster 20 neurons were highly distinct relative to other clusters in UMAP space (Fig. 2b), however we found that it was not entirely homogenous. While *Tcf4* was distributed evenly throughout cluster 20, one pole was enriched in *Pvalb* and the other pole was enriched in *Tac2* (Fig. 5a). Consistent with our scRNA-seq results, FISH analysis revealed extensive co-expression of *Tac2, Pvalb, Cplx1* and *Tcf4* with *Slc17a6* in the LM. Notably, *Pvalb* expression was limited to a subset of *Slc17a6*+ neurons in the LM (Fig. 5d). These data suggest that LM neurons comprise a transcriptionally-distinct population that appears to be confined to the anatomical boundaries of the LM.

**Figure 5:**
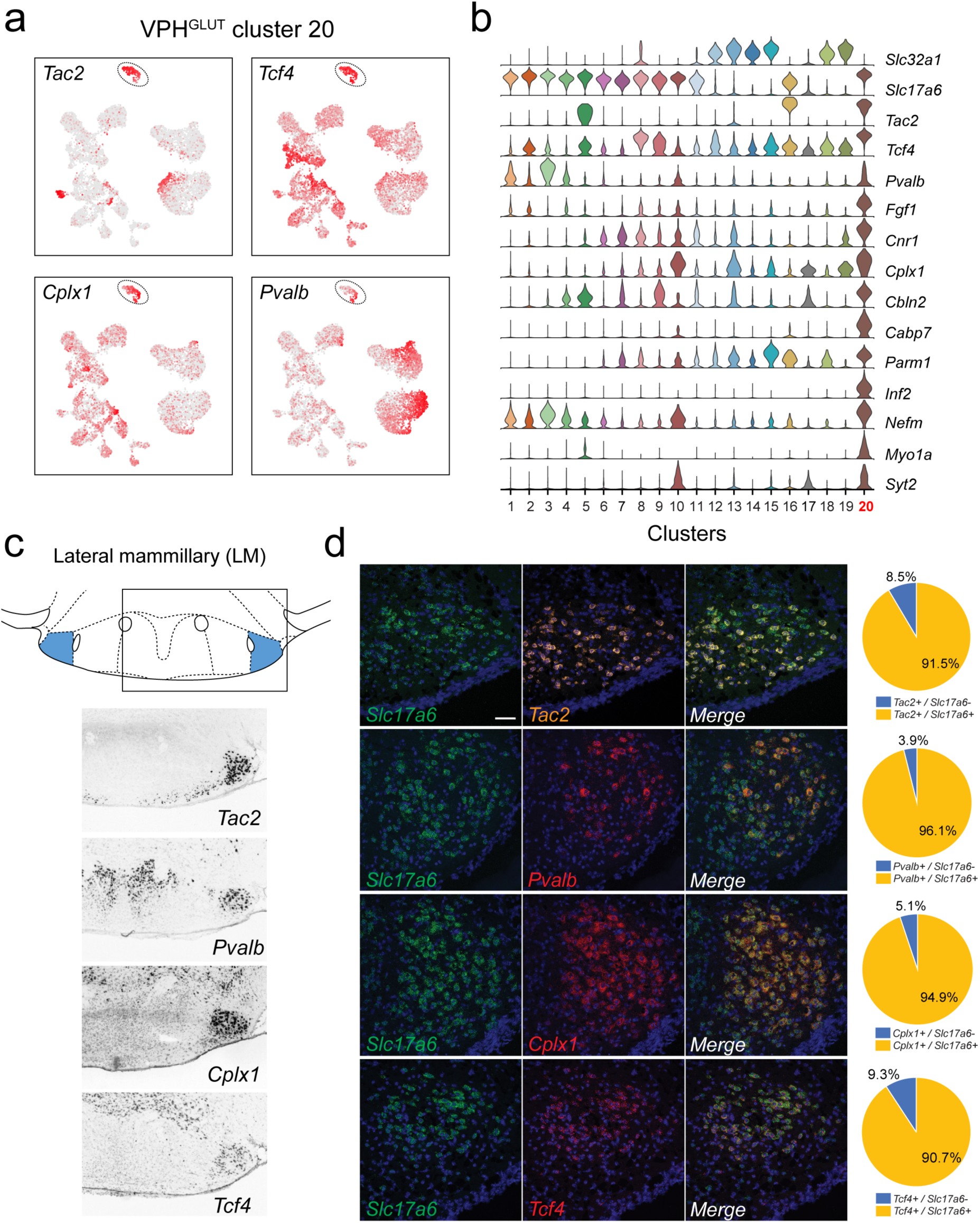
Identification of *Tac2*+ lateral mammillary (LM) neurons. **(a)** UMAP plots showing normalized expression of *Tac2, Tcf4, Cplx1* and *Pvalb* enriched in VPH^GLUT^ cluster 20 (circled). Violin plot showing discriminatory marker genes enriched in VPH^GLUT^ cluster 20 following *Slc32a1* and *Slc17a6* (top). **(c)** Mouse brain atlas schematic, modified from Paxinos and Franklin, 2012, showing the LM in a coronal section at distance from bregma of −2.92 mm (top). ISH images for *Tac2, Tcf4, Cplx1* and *Pvalb* from the ABA ^22^ (bottom). **(d)** Confocal micrographs (40x) of coronal sections of wild type mice and corresponding pie charts representing co-expression of mRNA for *Slc17a6* and *Tac2* (n=1210, 3 mice; upper), *Slc17a6* and *Pvalb* (n=1057, 3 mice, upper middle), *Slc17a6* and *Cplx1* (n=1269, 3 mice; lower middle), and *Slc17a6* and *Tcf4* (n=1642, 3 mice; lower), Scale bar (applicable to all micrographs) 50 μm.

### Multiple neuronal populations correspond to the medial mammillary region

We identified five VPH^GLUT^ clusters that appeared to be both closely interrelated to each other transcriptionally, and distinct from other neuronal clusters. VPH^GLUT^ clusters 1-5 were arranged in close proximity in UMAP space (Fig. 2b), with clusters 1 and 2 forming one contiguous cluster and clusters 3-5 forming another. Clusters 1-5 shared a number of common markers including genes that encode the neuropeptides *Cartpt, Cck, Adcyap1* as well as the transcription factor *Foxb1* (Fig. 6a and b). Cross-referencing *Cartpt, Foxb1, Cck* and *Adcyap1* with the ABA ^22^ indicate that VPH^GLUT^ clusters 1-5 correspond well to the medial mammillary region (Fig. 6c), which itself may be subdivided into cytoarchitecturally-defined anatomical compartments: median (MnM), medial (MM) and lateral (ML) subdivisions ^24^. Notably, *Cartpt, Foxb1* and *Cck*, show low expression in VPH^GLUT^ cluster 20 (LM) in our scRNA-seq data and correspondingly low expression the LM in ABA ISH data (Fig. 6c). In contrast, *Adcyap1* shows high expression in cluster 20 and high expression in the LM in ISH data, further reinforcing that clusters 1-5 represent the medial mammillary region, and cluster 20 represents the LM. Furthermore, other scRNA-seq analyses of whole mouse hypothalamus revealed a population of *Foxb1*+ neurons ascribed to the mammillary nuclei ^42,55^, while another scRNA-seq of the developing mouse hypothalamus. Multiplex FISH analysis revealed extensive co-expression of the neuropeptide transcripts *Cck, Adcyap1* and *Cartpt* with *Slc17a6* in the region (Fig. 6d). Other discriminatory markers that we found were common to clusters 1-5 include *Rprm, Cpne9, Ctxn3, Fam19a1* and *Gpr83* (Fig. 6b and Fig. S8a). Notably, we found that *Cpne9* (encoding copine-9) is uniquely enriched in both clusters 1-5 and cluster 20, as well as cluster 6 (Fig. S8a) suggesting that it may be a common marker for the entire MB, in addition to the PMd. Finally, we examined expression of other markers (*Pitx2, Lhx1, Lhx5, Cdh11, Sim1, Sim2* and *Nkx2.1*) that have previously been implicated in MB development ^55–65^and found significant alignment with the MB for virtually all markers (Fig. S8a).

**Figure 6:**
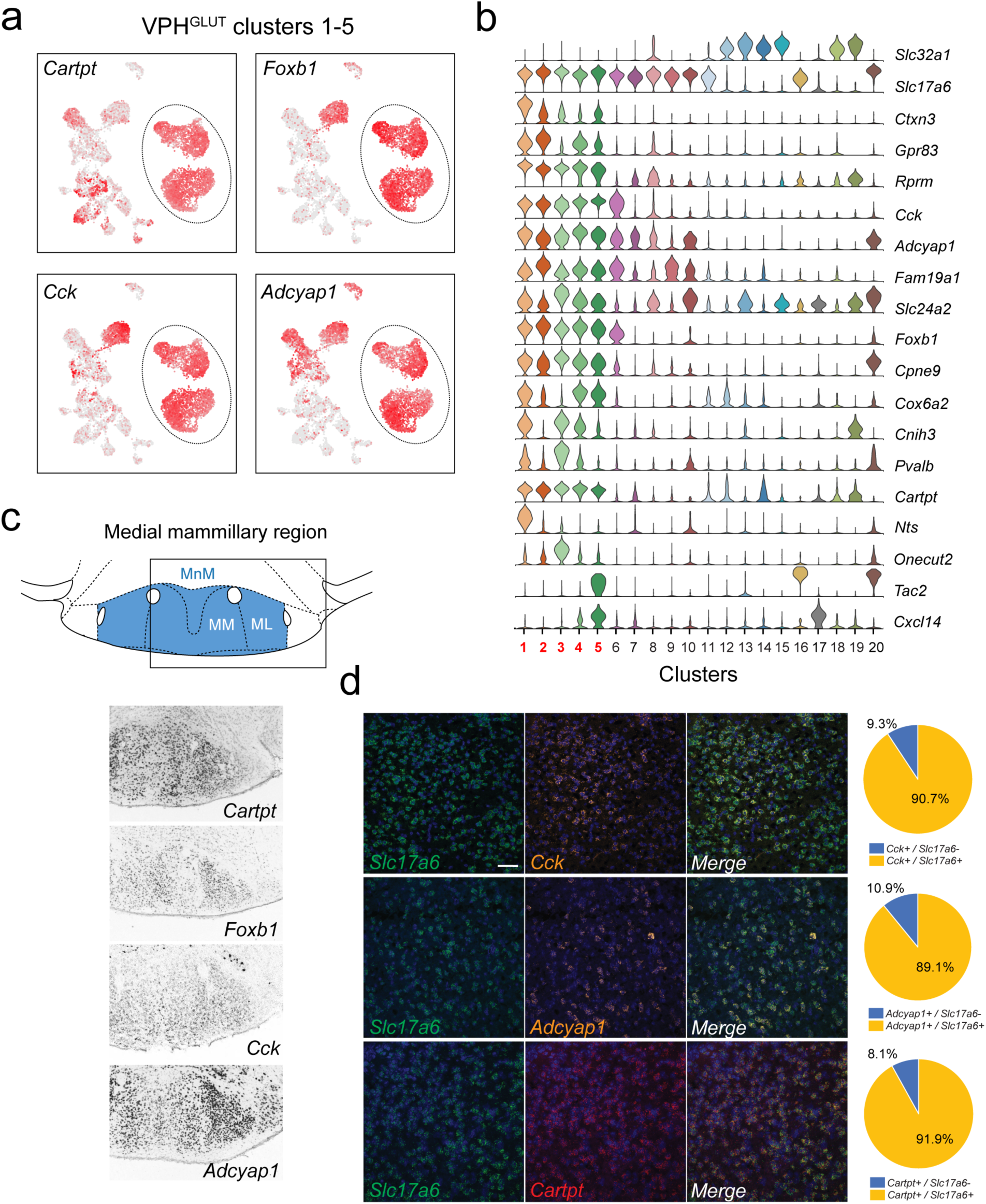
Identification of global markers for neurons in the medial mammillary region. **(a)** UMAP plots showing normalized expression of *Tac2, Tcf4, Cplx1* and *Pvalb* enriched in VPH^GLUT^ clusters 1-5 (circled). **(b)** Violin plot showing discriminatory marker genes enriched in VPH^GLUT^ clusters 1-5 following *Slc32a1* and *Slc17a6* (top). **(c)** Mouse brain atlas schematic, modified from Paxinos and Franklin, 2012, showing the medial mammillary, and its anatomical subdivisions (MnM, MM and ML) in a coronal section at distance from bregma of −2.92 mm (top). ISH images for *Cartpt, Foxb1, Cck* and *Adcyap1* from the ABA (Lein et al, 2007) (bottom) showing widespread expression throughout the medial mammillary. **(d)** Confocal micrographs (40x) of coronal sections of wild type mice and corresponding pie charts representing co-expression of mRNA for *Slc17a6* and *Cck* (n=2787, 3 mice; upper), *Slc17a6* and *Adcyap1* (n=1805, 3 mice; middle) and *Slc17a6* and *Cartpt* (n=2582, 3 mice; lower). Scale bar (applicable to all micrographs) 50 μm.

### Spatial segregation of transcriptionally-distinct medial mammillary subpopulations

We further investigated differentially expressed genes among VPH^GLUT^ clusters 1-5 (Fig. 7a), that comprise the medial mammillary region in an effort to determine if they exhibited any spatial organization with respect to the cytoarchitecurally-defined MnM, MM and ML subdivisions. We identified a suite of differentially-expressed genes that defined each of the transcriptionally distinct medial mammillary clusters 1-5 (Fig. 7b) with two discriminatory genes from each shown in UMAP plots (Fig. 7c). We went on to cross reference these makers with their spatial distribution within the medial mammillary region using the ABA ^22^) and found striking patterns of segmentation. Markers for cluster 1 neurons (*Nts*+, *Col25a1*+) appeared to be enriched in the MM subdivision, medial to the principle mammillary tract (pm), whereas markers for cluster 4 neurons (*Nos1*+, *Calb1*+) were found concentrated in the ML subdivision, lateral to the principle mammillary tract (pm) and medial to the fornix (f). Cluster 2 neurons (*Gpr83*+, *Spock3*+) were concentrated in the central portion of the MnM subdivision, whereas cluster 3 neurons (*Pvalb*+, *Slc24a2*+) were enriched in the more dorsal portion of the MnM. Interestingly, cluster 5 neurons (*Tac2*+, *Cxcl14*+) appeared to correspond to a thin rim of small diameter, *Tac2*+ neurons hugging the basal surface of the MM and ML (Fig. 7d), overlapping with the region typically referred to as the TMNv or VTM ^24^. These *Tac2*+ neurons may be distinguished from the larger diameter *Tac2*+ neurons confined to the LM (VPH^GLUT^ cluster 20). Despite being in the vicinity of TMNv HA neurons, 89.3% (176/197) of cluster 5 neurons were *Tac2*+ whereas only 2% (4/197) were *Hdc*+ indicating little to no overlap with HA neurons. In sum, we found that transcriptionally-distinct neuronal populations that comprise the medial mammillary region exhibit a remarkable degree of compartmentalization within the known anatomical subdivisions of the region, demarcated by the principle mammillary tract and fornix.

**Figure 7:**
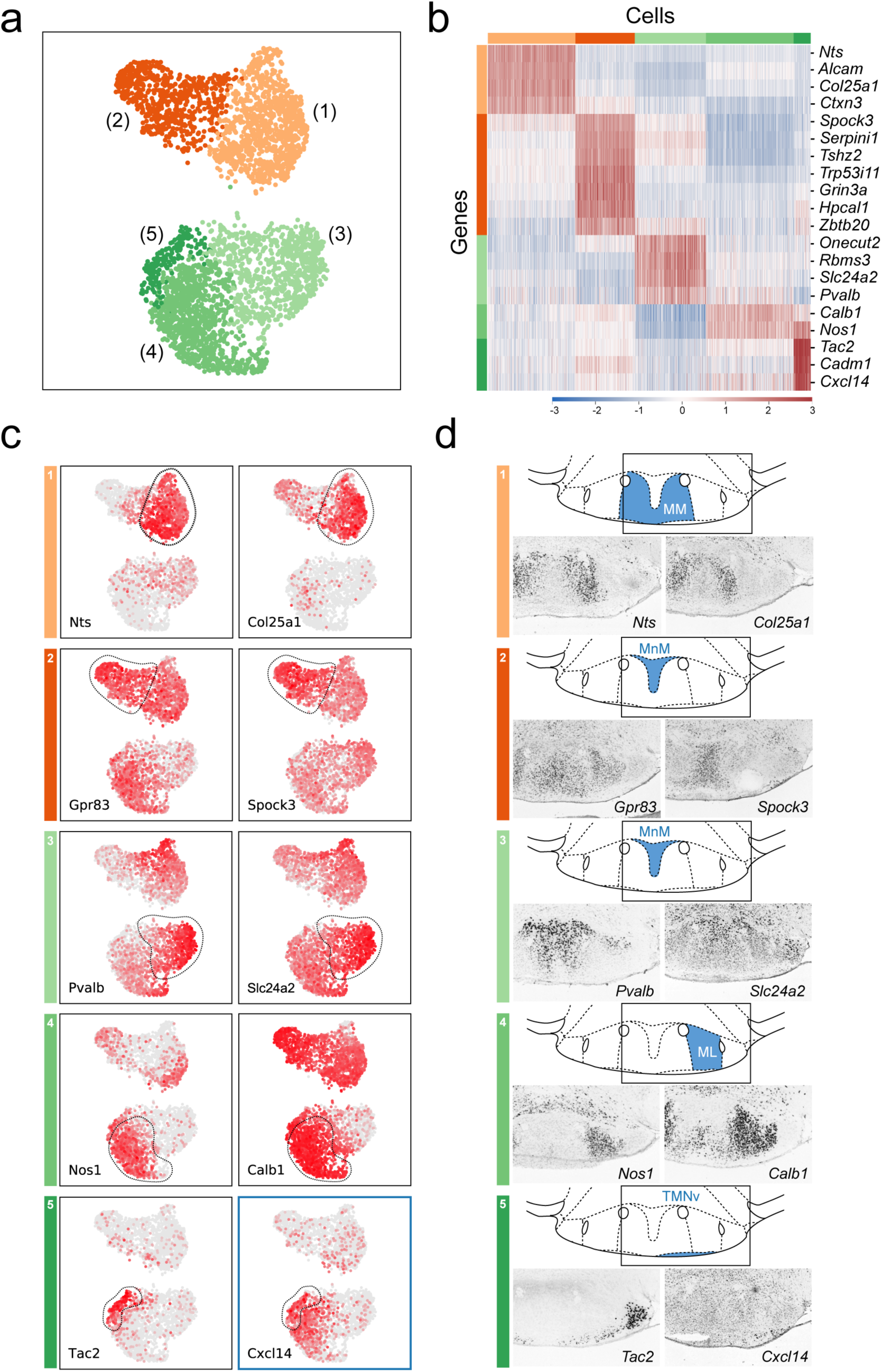
Anatomical compartmentalization of transcriptionally-distinct medial mammillary subpopulations. **(a)** UMAP plot showing just VPH^GLUT^ clusters 1-5 (detail of Fig. 2b). **(b)** Heatmap showing scaled expression of discriminatory genes across only VPH^GLUT^ clusters 1-5. Pairs of UMAP plots showing normalized expression of two discriminatory marker genes for each of VPH^GLUT^ clusters 1-5 (top to bottom): cluster 1 (*Nts, Col25a1*), cluster 2 (*Gpr83, Spock3*), cluster 3 (*Pvalb, Slc24a2*), cluster 4 (*Nos1, Calb1*) and cluster 5 (*Tac2, Cxcl14*). **(d)** Mouse brain atlas schematics, modified from Paxinos and Franklin, 2012, showing subregions of the medial mammillary in a coronal section at distance from bregma of −2.92 mm (top). ISH images for each of the two markers shown in (c) from the ABA ^22^ (bottom).

### Topographically-distinct projection targets of genetically-defined MM and ML neurons in the thalamus and midbrain

We next asked whether the cluster-specific discriminatory genes that we identified in our scRNA-seq analysis of the MB (summarized in Fig. 8a) may be used to genetically target specific anatomical subdivisions of the MB (summarized in Fig. 8b) and shed light on the organization of their projection targets. Neurons in the MB are known to make two highly specific projections in the brain: 1) a major, unidirectional projection to the anterior thalamic nuclei (ATN) through the mammillothalamic tract (mtt); and 2) a bidirectional connection to the midbrain ventral tegmental nucleus of Gudden (VTg), via the mamillotegmental tract (mtg) (summarized in Fig. 8c). The rodent ATN itself may be subdivided into three broad compartments, anteroventral (AV), anteromedial (AM) and anterodorsal (AD), each of which exhibit unique patterns of connectivity. In particular, the MM projects topographically to the AM, the ML to the AV and the LM to the AD ^7,66–68^. We therefore set out to test our prediction that the transcriptionally-distinct cell populations that we found to be segregated within specific subcompartments of the MB, exhibit a similar topographic mapping to the ATN. To this end, we used three cre recombinase mouse driver lines to target separate anatomical subregions of the MB by bilaterally injecting the MB with the viral anterograde tracer AAV-DIO-ChR2-EYFP (schematic in Fig. 8d). Using the *Slc17a6* (VGLUT2)-Cre driver line to broadly target VPH^GLUT^ neurons in the MB and their projections, we found labeling throughout the medial mammillary region at the injection site (Fig. 8e), but largely sparing the LM, shown in a representative mouse. In addition to finding a high density of labeled axons in the mtt and mtg, we observed a plexus of fibers in the VTg and a high density of fibers in much of the ATN (AV and AM, but not AD, consistent with the latter being the target of LM projections). Next, using the *Nts*-Cre line to selectively target VPH^GLUT^ cluster 1 neurons, that we had previously mapped to the MM subdivision, we found labeling at the injection site between the MM and ML and labeled axons in the mtt, mtg and VTg, shown in a representative mouse. However, in the ATN, the AM subdivision appeared to be selectively targeted, while sparing the AV and AD (Fig. 8e). Finally, using the *Calb1*-Cre driver to target VPH^GLUT^ cluster 4 neurons, that we had mapped to the ML subdivision, we found clear labeling in the ML at the injection site (with some weak labeling in the MM/MnM) and labeled axons in the mtt, mtg and VTg, shown in a representative mouse. Interestingly, and in stark contrast to our *Nts*-cre results, we observed highly specific labeling of the AV subdivision of the ATN, while sparing the AM and AD. These data are consistent with immunohistochemical evidence in guinea pig suggesting that CART (encoded by *Cartpt*) and calbindin (encoded by *Calb1*) are co-localized in the ML region of the MB and calbindin-immunoreactive fibers are concentrated in the AV subdivision of the ATN ^69^. Taken together, these anterograde tracing data suggest that not only are molecularly-defined subpopulations of MB neurons highly segregated within the anatomical subdivisions of the MB (MnM, MM, ML and LM), but these subpopulations appear to project to distinct domains within the ATN, consistent with previous anterograde and retrograde tracing data without genetic specificity ^7,66–68^.

**Figure 8:**
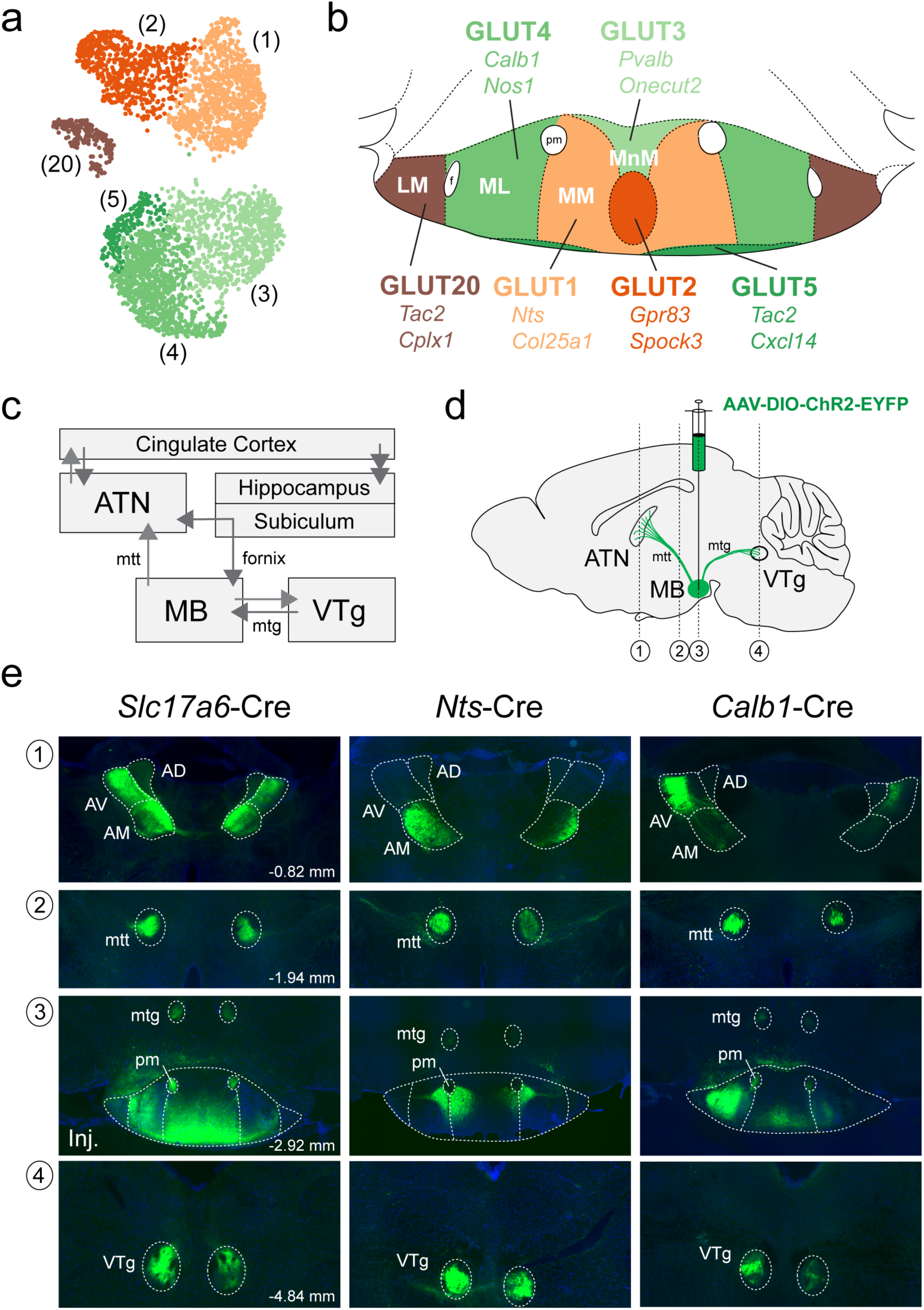
Anterograde projections of transcriptionally-distinct medial mammillary subpopulations to the anterior thalamus and midbrain. **(a)** UMAP plot showing just VPH^GLUT^ clusters 1-5 and 20 (details from Fig. 2b). **(b)** Mouse brain atlas schematic modified from Paxinos and Franklin, 2012, showing subregions of the MB in a coronal section at distance from bregma of −2.92 mm. Color-coding of subdivisions indicate correlations between transcriptionally distinct subpopulations in (a) and anatomical subdivisions within the MB: cluster 1 (MM, light orange), cluster 2 (ventromedial MnM, dark orange), cluster 3 (dorsomedial MnM, light green), cluster 4 (ML, medium green), cluster 5 (TMNv or ventrobasal MM/ML, dark green) and cluster 20 (LM, brown). **(c)** Schematic of connections between the MB, hippocampal formation, ATN and VTg comprising the Papez circuit. **(d)** Schematic parasagittal section illustrating anterograde viral tracing using AAV-DIO-ChR2-EYFP injection in the MB of three cre driver lines (*Slc17a6*-Cre, *Nts*-Cre and *Calb1*-Cre). Number labels indicate imaged coronal sections. **(e)** Representative fluorescence images of coronal sections for anterograde tracing in *Slc17a6*-Cre (n=3 mice; left), *Nts*-Cre (n=3 mice; middle) and *Calb1*-Cre (n=2; right) mice at the following approximate distances from bregma: 1) ATN, −0.82 mm; 2) mtt, −1.94 mm; 3) MB injection site, −2.92 mm; and 4) VTg, −4.84 mm. All sections were counterstained with DAPI (blue). Abbreviations: AV (anteroventral), AM (anteromedial), AD (anterodorsal), mtt (mammilothalamic tract), mtg (mammillotegmental tract), pm (principle mammillary tract).

## DISCUSSION

In this work, we performed a systematic single cell transcriptomic census of the molecular and spatial organization of cell types in the mouse VPH. Using droplet-based scRNA-seq of >16,000 cells, we identified the molecular markers that define 18 non-neuronal and 20 neuronal cell clusters. We identified a diversity of mostly novel, and some previously identified, neuronal cell populations, many of which could be mapped back to clearly defined anatomical compartments within the VPH (including the Arc, TMN, SUM, PMd, PMv and approximately six subdivisions of the MB including the MnM, MM, ML and LM), using both ISH data from the ABA and multiplexed FISH. In particular, we observed that transcriptionally-distinct MB neurons are both confined to anatomically segregated compartments and, using three genetically-targeted neuronal populations in the MB, appear to project to precise anatomical targets in the ATN. This molecular census provides a rich resource for interrogating genetically-defined VPH circuits and their myriad roles in physiology and behavior.

In our unsupervised analysis of VPH neuronal populations, numerous molecularly distinct clusters emerged (Fig. 2), but a broader dichotomy could be found based on the expression of genes necessary for the synthesis and vesicular release of GABA and glutamate.

The primary discriminatory genes in this regard are the largely mutually exclusive expression of *Slc32a1* (VGAT) and *Slc17a6* (VGLUT2) resulting in our identification of 13 populations of nominally glutamatergic neurons and 6 populations of nominally GABAergic neurons. Overall, the broad division between nominally GABAergic and glutamatergic populations aligns with observations from other recent scRNA-seq analyses of whole mouse brain ^26,27^, whole hypothalamus ^42^ and hypothalamic subregions ^43,44,70–73^. One notable exception is VPH^HA^ cluster 17 which expresses low levels of both *Slc32a1* and *Slc17a6*, and we identified as having a histaminergic phenotype (Figs. 2 and 4). Another notable exception is a dual identity cluster 8 subcluster, which exhibits high co-expression of both *Slc32a1* and *Slc17a6*, and corresponds well to markers of the SUM (Fig. S6) and, possibly, to a known dual phenotype, hippocampal-projecting population previously identified in the SUM ^36–39^.

Within each fast neurotransmitter identity, discrete classifications of VPH neurons were arrived at based on a suite of other discriminatory markers with few classifications being made on the basis of single makers alone. Most neuronal clusters are also defined by neuropeptide transcripts, which are associated with both well-known and novel populations. The co-expression of both fast neurotransmitter and multiple neuropeptide markers in single neurons is suggestive of a capacity for co-transmission ^74^. Other markers invariably include some combination of transcripts that encode transcription factors, calcium binding proteins, synaptic and extracellular matrix proteins, receptors and other gene categories. That these combinations of markers specify the identity of distinct populations of VPH neurons likely reflects a convergence of their unique developmental history, neurochemistry and specific synaptic connectivity within functional circuits.

While several of the diverse neuronal clusters that emerged from our scRNA-seq analysis were readily identifiable based on well-known signatures (ex. *Kiss1*+/*Tac2*+/*Pdyn*+ KNDy Arc neurons, *Agrp*+/*Npy*+ Arc neurons, *Hdc*+ TMN HA neurons, etc.), most were novel and differentially expressed genes required both cross-referencing with the ABA ^22^ and extensive multiplexed FISH to effectively trace them to anatomical regions within the VPH. We found that a number of novel neuronal cell populations exhibited striking compartmentalization within distinct VPH nuclei. This was especially apparent in our mapping of VPH^GLUT^ clusters 1-5 and 20 to subcompartments within the MB. The MB is a conserved diencephalic structure across rodents, primate models and humans and represents an important link between subicular outputs of the hippocampal formation (through the fornix) and the anterior thalamus (through the mtt), which itself has reciprocal projections with the cingulate/retrosplenial cortices. MB neurons also make bidirectional connections with the midbrain VTg (through the mtg) ^5–9,67,68^. MB neurons, as a key component of this limbic pathway, are critical for spatial memory as lesions to the MB and its outputs through the mtt and mtg, result in severe deficits in spatial memory in animal models and human clinical data ^5,6,8,9,68,75–77^. For example, Korsakoff’s syndrome, caused by thiamine deficiency associated with chronic alcoholism, is characterized by selective degeneration of the MB and is accompanied by severe amnesia ^78,79^.

Consistent with our identification of VPH^GLUT^ clusters 1-5 as representing the medial mammillary region, several of the discriminatory marker genes that emerged from our analysis result in spatial memory deficits when mutated. For example, *Foxb1* is well-known to define the MB during early embryonic development ^55,61,80–82^ and *Foxb1*-null mice exhibit dysgenesis of both the MB and mtt ^81–83^ and display defects in spatial memory tasks, particularly spatial navigation and working memory ^82^. These data suggest that the MB may have specific roles to play in spatial memory processing. Furthermore, the enrichment of the recently deorphanized receptor *Gpr83* ^84^ that we observed in this region is aligned with previous anatomical work showing high expression in the MB ^85^ and behavioral experiments showing that *Gpr83*-null mice also exhibit impaired spatial learning ^86^. These data indicate that VPH^GLUT^ clusters 1-5 share a suite of common markers that align well with the known development and function of the medial mammillary region of the MB.

Interestingly, a selection of the defining marker genes for the medial mammillary region (clusters 1-5) (Figs. 6, 7 and S8), were also found to be enriched in some combination of other clusters we ascribed to the LM (cluster 20), PMd (cluster 6), PMv (cluster 7), SUM (cluster 8) and others. For example, we found that while *Cck, Foxb1, Pitx2, Lhx1, Lhx5, Sim1* were indeed all enriched in clusters 1-5, they were all additionally expressed in cluster 6, among others. In addition to the MB, *Foxb1* is known to also be expressed in the developing PMd ^81,83^, which is linked anatomically and functionally with the MB, including a parallel projection to the ATN ^10,83,87^. Furthermore, *Foxb1*-null mice not only exhibit defects in spatial memory ^82^ but abnormal nurturing behavior including pup retrieval and nest-building ^81^. These data suggest a possible defect in PMv-associated reproductive behavior ^11^ and/or PMd-associated threat-response or defensive behavior ^10,88,89^. Taken together, our transcriptomic data reinforce the notion that the MB, PMd, PMv and perhaps other surrounding VPH structures, share an anatomical and functional interrelationship, rooted in their shared developmental history ^57,65^.

In mapping differentially expressed markers of VPH^GLUT^ clusters 1-5 and 20 to the MB, we found that the MB is not only more transcriptionally diverse than previously appreciated, but distinct MB neurons also exhibit a remarkable degree of segregation within long-established, anatomically-defined subdivisions of the MB. While the primary anatomical division in the MB is between medial and lateral, the medial mammillary may be further subdivided into multiple subcompartments (MnM, MM and ML) on the basis of Nissl and Golgi staining, cell morphology, white matter boundaries (pm, fornix, etc) and patterns of connectivity ^24,67,68,90,91^. However, cell type diversity and the spatial organization of cell types within the MB is poorly understood. Although a recent scRNA-seq analysis of the whole hypothalamus appeared to capture the VPH, it resolved only a single population of *Foxb1*+/*Cartpt*+ neurons assigned to the MB ^42^. In contrast, we resolved at least six distinct populations that we subsequently mapped to six anatomical subdivisions of the MB (summarized in Fig. 8a and b). For example, VPH^GLUT^ cluster 20 (*Tac2*+/*Cplx1*+) was confined to the LM and was distinct from cluster 4 (*Nos1*+/*Calb1*+), which was confined to the ML, which itself was distinct from cluster 1 (*Nts*+/*Col25a1*+), which broadly corresponded to the MM. The patterns of discriminatory genes for clusters 2 (*Gpr83*+/*Spock3*+) and cluster 3 (*Pvalb*+/*Slc24a2*+) were more complex. Cluster 3 markers appeared to correspond to the more dorsal MnM, whereas cluster 2 markers were more ventromedially located, bounded by the MM on either side, hence the slightly modified boundaries schematized in Fig. 8b. Cluster 5 (*Tac2*+/*Cxcl14*+) appeared to correspond to a thin rim of neurons on the ventrobasal surface of the MB and distinct from *Tac2*+ neurons in the LM. This pattern of cell type segmentation was striking, especially juxtaposed with the highly diffuse spatial organization of transcriptionally-distinct neuronal populations in the adjacent LHA ^44^.

The significance of this highly compartmentalized pattern of gene expression may lie in the specific connectivity between MB subcompartments and other regions of the brain. The importance of MB projections to the ATN and VTg is highlighted by previous work showing that specific lesions to the mtt and VTg, but not to the postcommissural fornix, result in deficits in spatial memory, suggesting that MB-ATN and MB-VTg connectivity have critical roles in memory, likely independent of hippocampal input ^76^. We found that genetically-defined MB neurons, using markers we identified in our scRNA-seq analysis to specify MB subpopulations, project to distinct thalamic and midbrain targets. In particular, while *Slc17a6*+ MB neurons project to the AV and AM, *Nts*+ MM neurons project specifically to the AM, while *Calb1*+ ML neurons project specifically to the AV. All three MB projections target the VTg while none targeted the AD. Our genetically-targeted anterograde tracing is consistent with the known topography of MB-ATN projections ^7,66–68^ and suggests that the spatial architecture of these circuits may be explained by an underlying molecular organization. Furthermore, the AM, AV and AD subregions of the ATN each have differential patterns of reciprocal connectivity with different cortical regions. For example, aside from reciprocal connections with the anterior cingulate, the AM is uniquely interconnected with the prelimbic cortex ^7,67^. Aside from topographic ATN projections, the spatial organization of MB circuits is also likely a reflection of topographic projections from specific subicular subregions/cell types ^91–93^, which also exhibit a broader molecular and spatial organization ^94^. Future work combining transcriptomic profiling, functional connectomics, manipulation and monitoring of genetically-defined neurons in the subiculum, MB, VTg, ATN and cingulate/retrosplenial cortices may reveal an overarching organization of parallel circuits within this brain-wide memory system. Recent large-scale efforts towards molecular profiling and detailed connectivity analysis among other thalamic pathways, have revealed novel insights into the overarching logic of thalamic organization ^95,96^. Taken together, our transcriptomic analysis of MB subpopulations may shed light on the construction of MB circuits during development, their differential connectivity, regulation of excitability, repertoire of synaptic signaling mechanisms and specific functional roles in spatial memory.

Finally, given the important and conserved role of the MB as a node in a brain-wide memory system and multiple aspects of spatial and episodic memory, it is significant that the MB may have a key role in the pathogenesis of Alzheimer’s Disease (AD). Evidence from imaging of AD patients and postmortem examination of AD-afflicted brains suggest that the MB, fornix and mtt all show varying degrees of atrophy or degeneration that may be correlated with cognitive decline and memory loss ^97–100^. Recent work has implicated the MB and other components of the Papez circuit as sites of particular vulnerability in the pathogenesis of AD in mice ^101^. Using a quantitative measure of beta-amyloid deposition in the whole brain of 5XFAD mice, a mouse model of AD ^102^, the MB was identified as among the earliest sites of beta-amyloid deposition in the brain, appearing as early as two months after birth. This observation was accompanied by an increase in medial mammillary neuron excitability, that when chemogenetically silenced, resulted in reduced beta amyloid deposition ^101^. In addition, other VPH subpopulations may exhibit selective vulnerability in AD as evidenced by the presence of AD pathological features in the TMN and dramatic loss of TMN HA neurons in *post mortem* AD-afflicted brains ^103–106^, likely contributing to sleep-wake disturbances commonly observed in AD patients ^107^. Taken together, these clinical observations and intriguing evidence from disease models underscore the critical need to elucidate both the basic biology of the MB and other VPH cell types, the circuits they give rise to and their potential vulnerability in the early stages of AD pathogenesis.

Overall, our analysis of the molecular and spatial organization of VPH cell types provides the foundation for a more detailed understanding of the cellular composition and wiring diagram of the VPH. In particular, our analysis reveals the molecular underpinnings of the modular organization of both the MB and its topographic projections to subregions of the anterior thalamus as well as the midbrain. In addition, this work serves as a rich resource for cell type-specific deconstruction of VPH circuit function, through pharmacological tools and the use of genetically-targeted manipulation and monitoring methodology. Overall, our VPH cell type census, along with other recent hypothalamic scRNA-seq analyses ^26,42–44,55,70–73^, contributes to a more comprehensive picture of the cell type diversity and spatial organization of the mammalian hypothalamus, as well as informing the molecular, cellular and synaptic mechanisms of hypothalamic circuit function in health and disease.

## MATERIALS AND METHODS

### Ethics statement

All experiments were performed in accordance with the ethical guidelines described in the National Institutes of Health Guide for the Care and Use of Laboratory Animals and were approved by the Institutional Animal Care and Use Committee of the University of Connecticut and of the Jackson Laboratory for Genomic Medicine.

### Animals

To collect VPH neurons scRNA-seq analysis as well as fluorescence *in situ* hybridization experiments, we used both male and female C57BL/6 (JAX stock #000664) mice. For anterograde tracing experiments, we used the following cre recombinase driver lines: 1) *Slc17a6*^tm2(cre)Lowl^/J knockin mutant mice (JAX stock #016963, referred to here as *Slc17a6*-Cre mice) ^108^. 2) *Nts*-Cre knock-in mutant mice (B6;129-Nts^tm1(cre)Mgmj^/J) (JAX stock # 017525) ^109^. 3) *Calb1*-IRES2-Cre-D knock-in mutant mice (B6;129S-Calb1^tm2.1(cre)Hze^/J) (JAX stock # 028532, referred to here as *Calb1*-Cre). All mice were fed *ad libitum* and kept on a 12 h light-dark cycle.

### scRNA-seq cell capture and sequencing

Hypothalamic brain slices were obtained from juvenile (p30-34) mice, from a total of 5 male mice and two female mice over two independent harvests. The first harvest consisted of 3 male (pooled), 3 female (pooled), while the second consisted of 2 males (pooled) and 2 females (pooled), following previously described procedures ^44^. Briefly, mice were anesthetized with isoflurane, then rapidly sacrificed by decapitation during the same time period (morning, 09:00-11:00). Brain slices were cut using a vibrating microtome (Campden Instruments) at a thickness of 225 μm in ice cold high-sucrose slicing solution consisting of the following components (in mM): 87 NaCl, 75 sucrose, 25 glucose, 25 NaHCO_3_ 1.25 NaH_2_PO_4_, 2.5 KCl, 7.5 MgCl_2_, 0.5 CaCl_2_ and 5 ascorbic acid saturated with 95% O_2_/5% CO_2_. Slices were then enzyme treated for ∼15 min at 34°C with protease XXIII (2.5 mg/mL; Sigma) in a high-sucrose dissociation solution containing the following (in mM): 185 sucrose, 10 glucose, 30 Na_2_SO_4_, 2 K_2_SO_4_, 10 HEPES buffer, 0.5 CaCl_2_, 6 MgCl_2_, 5 ascorbic acid (pH 7.4) and 320 mOsm. Slices were washed three times with cold dissociation solution then transferred to a trypsin inhibitor/ bovine serum albumin (TI/BSA) solution (10 mg/mL; Sigma) in cold dissociation solution. Four to five slices were obtained from each animal that approximately corresponded to mouse brain atlas images representing the distance from bregma −2.54, −2.70, −2.92 and −3.16 mm which includes the VPH. The VPH was dissected from each slice using a fine scalpel and forceps under a dissecting microscope and each slice was imaged and subsequently mapped onto mouse atlas images (Fig 1b) ^24^. Microdissected tissue was kept in cold TI/BSA dissociation solution until trituration. Immediately before dissociation tissue was incubated for ∼10 min at 37°C, then triturated with a series of small bore fire-polished Pasteur pipettes. Single cell suspensions were passed through a 40 μm nylon mesh filter to remove any large debris or cell aggregates and kept on ice until single-cell capture.

Cell viability for each sample was assessed on a Countess II automated cell counter (ThermoFisher), and up to 12,000 cells were loaded for capture onto an individual lane of a Chromium Controller (10X Genomics). Single cell capture, barcoding and library preparation were performed using the 10X Chromium platform ^23^ according to the manufacturer’s protocol (#CG00052) using version 2 (V2) chemistry for the first set of male and female samples and version 3 (V3) chemistry for the second set. cDNA and libraries were checked for quality on Agilent 4200 Tapestation, quantified by KAPA qPCR. The two V2 chemistry libraries were sequenced on individual lanes of an Illumina HiSeq4000 and the V3 chemistry libraries were pooled at 16.67% of lane of an Illumina NovaSeq 6000 S2 flow cell each, all samples targeting 6000 barcoded cells with an average sequencing depth of 50,000 reads per cell.

### scRNA-seq data processing, quality control, and analysis

Illumina base call files for all libraries were converted to FASTQs using bcl2fastq v2.20.0.422 (Illumina) and FASTQ files were aligned to the mm10 (GRCh38.93, 10X Genomics mm10 reference 3.0.0) using the version 3.0.2 Cell Ranger count pipeline (10X Genomics), resulting in four gene-by-cell digital count matrices. Downstream analysis was performed using Scanpy (v1.3.7) ^110^. Initial quality control was performed on each library individually and cells were excluded from downstream analysis by the following criteria: fewer than 2,000 transcripts, fewer than 1,000 genes, more than 50 hemoglobin transcripts, and more than 15% mtRNA content. Genes with fewer than 3 counts in at least 3 cells were also excluded from downstream analysis. These filtering criteria resulted in a substantial increase in dataset quality but a dramatic reduction in called cells; the resulting individual counts matrices were reduced to 5,191 and 3,223 cells for the male and female V2 libraries, respectively, and 4,622 and 3,955 cells for the male and female V3 libraries. These matrices were then concatenated together, resulting in an initial aggregated counts matrix of 16,991 cells by 20,202 genes. This aggregated counts matrix was normalized by the total number of counts per cell then multiplied by the median number of counts across all cells and finally log_2_ transformed.

The 1,500 most highly variable genes as measured by dispersion were selected for the computation of principal components (PCs). Prior to the computation of PCs, mitochondrial, ribosomal, hemoglobin, cell cycle (homologs of those defined in Macosko et al, 2015 ^111^) genes, and the specific genes *Xist, Fos, Fosb, Jun, Junb*, and *Jund* were excluded from this set highly variable genes. The first 50 PCs were computed and subsequently corrected for batch effects related to 10X chemistry version and mouse sex using Harmony ^112^ (with theta_chemistry = 2, theta_sex = 0.5). Without this batch correction, clusters were stratified by 10X chemistry (shown in Fig. S1). The 15 PCs with highest variance ratios were used to construct a k=15 nearest-neighbor graph (k-NN graph, where distance is measured using cosine distance) and a 2D UMAP embedding was generated using this graph (min_dist = 0.5). Initial clusters were assigned via the Leiden community detection algorithm on this k-NN graph, resulting in 31 initial clusters.

Marker genes for each cluster, computed by a “one-versus-rest” methodology comparing mean expression of every gene within a cluster to the expression in all other cells, were used to assign putative cell types to each cluster. Of the 31 initial clusters, 8 clearly exhibited signatures of two cells types and 518 cells from these 8 clusters were excluded from further analysis. The remaining cells were classified as neuronal or non-neuronal using a simple two state Gaussian mixture model. Briefly, the median expression of *Snap25, Syp, Tubb3, and Elavl2* in each cluster was used to fit the model and classify the clusters as neuronal (high expression) and non-neuronal (low expression). We also tested classifying each individual cell using this model which lead to qualitatively equivalent classification. The cells for each classification were subsequently reanalyzed separately.

Neuronal cells underwent another round of filtering, with fewer than 3,500 transcripts and 2,200 genes being excluded. The expression matrix of the remaining 9,304 cells was normalized, batch corrected, and embedded with UMAP as described above with the only alteration being 20 PCs were used to build a k=10 NN graph. Clustering with Leiden community detection led to 20 neuronal clusters. Non-neuronal cells were reanalyzed without additional filtering, and the expression matrix for the 6,069 non-neuronal cells was reanalyzed as described for the neuronal cells; Leiden clustering yielded 18 non-neuronal clusters.

### Fluorescent *in situ* hybridization

FISH was carried out as previously described ^44^ Briefly, to prepare tissue sections for FISH, male juvenile wild type C57BL/6 mice (P30–47) were anesthetized with isoflurane, decapitated and brains were dissected out into ice-cold sucrose. Brains were frozen on dry ice and cryosectioned at a thickness of 14 µm directly onto SuperFrost Plus microscope slides. Sections were then fixed with 4% paraformaldehyde (PFA) at 4 °C for 15 min, and dehydrated for 5 mins each in 50, 70 and 100% ethanol. RNAscope 2.5 Assay (Advanced Cell Diagnostics) was used for all FISH experiments according to the manufacturer’s protocols. All RNAscope FISH probes were designed and validated by Advanced Cell Diagnostics. ISH images from the Allen Brain Institute were acquired from the publically available resource, the Allen Mouse Brain Atlas (www.mouse.brain-map.org/) ^22^ and were acquired with minor contrast adjustments and converted to grayscale to maintain image consistency.

### Anterograde viral tract-tracing

For anterograde tracing experiments, male P30-42 *Slc17a6*-cre, *Nts*-Cre and *Calb1*-Cre mice were bilaterally injected with 50-100nL of AAV2-Ef1α-DIO-hChR2(H134R)-EYFP (UNC Viral Core, Diesseroth Lab) into the MB (anteroposterior (AP): - 2.90 mm, mediolateral (ML): ±0.04 mm, dorsoventral (DV): −5.20 mm) and allowed to incubate for 4-6 weeks. *Slc17a6*-cre mice (n=3) were injected with 100 nL while *Nts*-Cre mice (n=3) and *Calb1*-cre mice (n=2) were injected with 50 nL. For histology, mice were anesthetized with ketamine/xylazine and transcardially perfused with 10 mL of 0.125 M saline followed by 40 mL of 4% PFA in 1X PBS (pH 7.4). Brains were then dissected and post-fixed for 24 hrs in 4% PFA, followed by cryoprotection in 30% sucrose for 48 hrs. Brains were flash frozen with cold isopentane and stored at −80°C. Frozen brains were cut into 50 μm thick coronal sections containing the ATN, LHA, MB and VTg on a cryostat (Leica 3050s) into 1X PBS. Slices were then mounted onto slides with Vectashield Hardset mounting media containing DAPI (Vector Labs). Regions of interest were imaged at 10X magnification on a fluorescence microscope (Keyence BZ-X700). Images were processed with ImageJ, Adobe Photoshop CS and Adobe Illustrator CC. Based on *post hoc* histological evaluation of the injection site, mice were excluded from the behavioral analysis if MB injection sites were absent or off-target.

### Data and code availability

Raw sequencing data and counts matrices are available in the Gene Expression Omnibus, accession number GSE146692. We further provide the final analyzed datasets in the form of H5AD ^110^ and Loupe Cell Browser data files, as well as all code to reproduce the analysis and figure panels, on GitHub: https://github.com/TheJacksonLaboratory/ventroposterior-hypothalamus-scrna-seq.

## ACKNOWLEDGEMENTS

We gratefully acknowledge R. Kanadia, A. Tzingounis, for helpful discussions and comments on the manuscript, and M. Picciotto for helpful discussions. We also gratefully acknowledge D. Luo for single cell capture and scRNA-seq library preparation, A. Mosley for genotyping support, C. O’Connell for imaging support. We would also like to thank A. Fujita and Y. Huang, as well as all members of the Jackson lab for support, assistance and helpful discussions. Project supported by the National Institutes of Health grant R01MH112739 (to A.C.J.), a Connecticut Innovations Regenerative Medicine Research Fund grant 15-RMD-UCHC-01 (to PR) and an NIH Shared Instrumentation Grant S10OD016435 (to A. Nishiyama) for imaging support.

## AUTHOR CONTRIBUTIONS

L.E.M. designed experiments, performed microdissections, single cell isolation, FISH experiments/analysis, stereotactic injections, neuroanatomical tracing/imaging and edited the manuscript. W.F. developed bioinformatics pipelines, performed bioinformatics analysis and edited the manuscript. K.S, L.W. and E.B performed sectioning, mounting and neuroanatomical tracing/imaging. M.B performed bioinformatics analysis on an early iteration of the dataset. P.R. designed experiments, provided intellectual guidance on bioinformatics and edited the manuscript. A.C.J. conceived and supervised the study, designed experiments, performed analysis and wrote the manuscript.

## COMPETING INTERESTS

The authors declare no competing interests.

## SUPPLEMENTARY FIGURES

**Figure S1:**
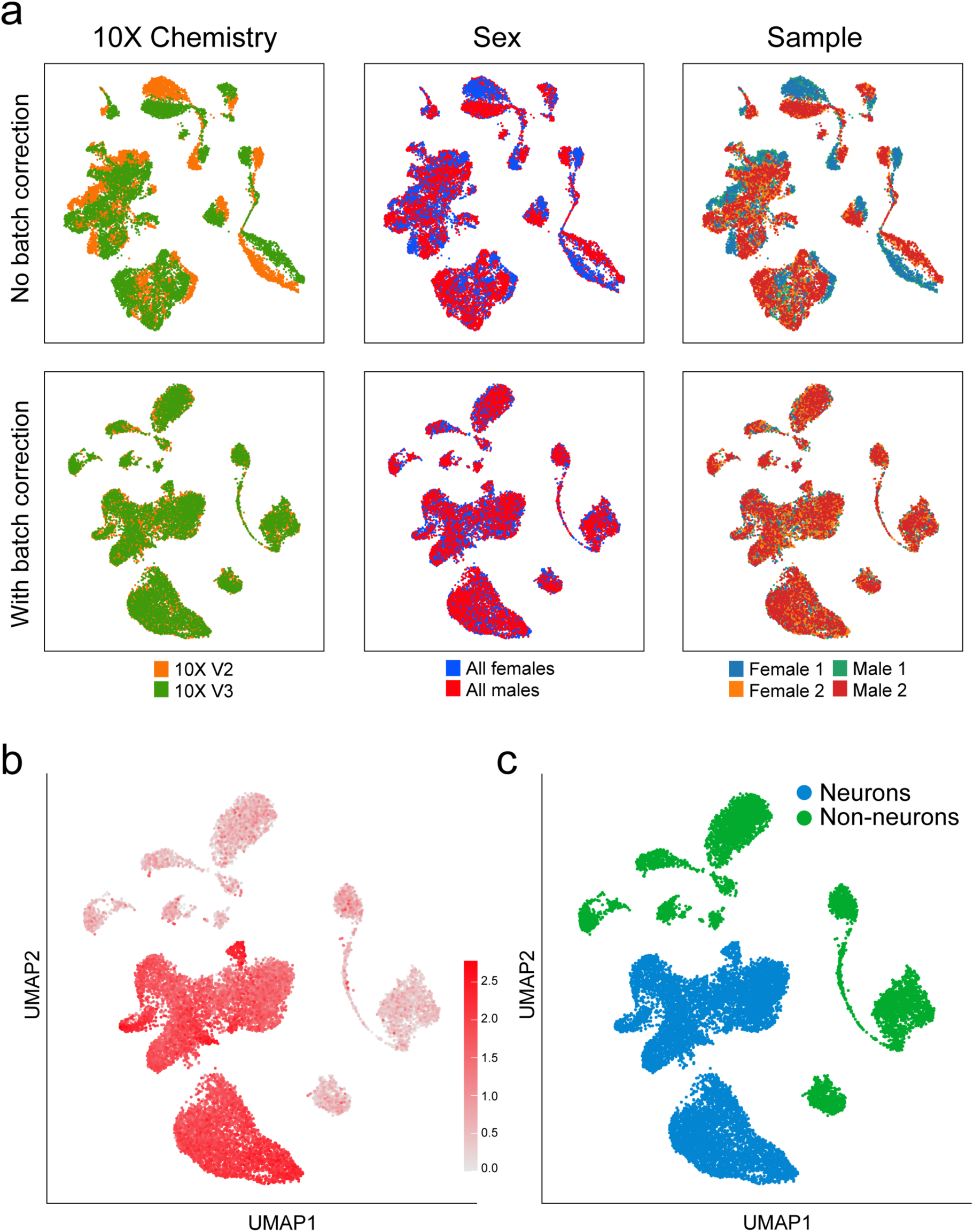
Batch correction for sex and 10X Genomics chemistry versions. **(a)** When libraries were combined bioinformatically, we assessed the need for batch correction by visualizing the libraries with (lower) and without (upper) Harmony batch correction ^112^. Batch effects correlated with 10X Genomics chemistry version were observed, but no batch effects were associated with mouse sex. **(b)** UMAP plot of average normalized expression of pan-neuronal markers *Snap25, Syp, Tubb3* and *Elavl2* across all cells prior to the first iteration of unsupervised clustering. **(c)** A two-class Gaussian mixture model was trained using the expression of these four genes to segregate neuronal cells (blue) from non-neuronal cells (green).

**Figure S2:**
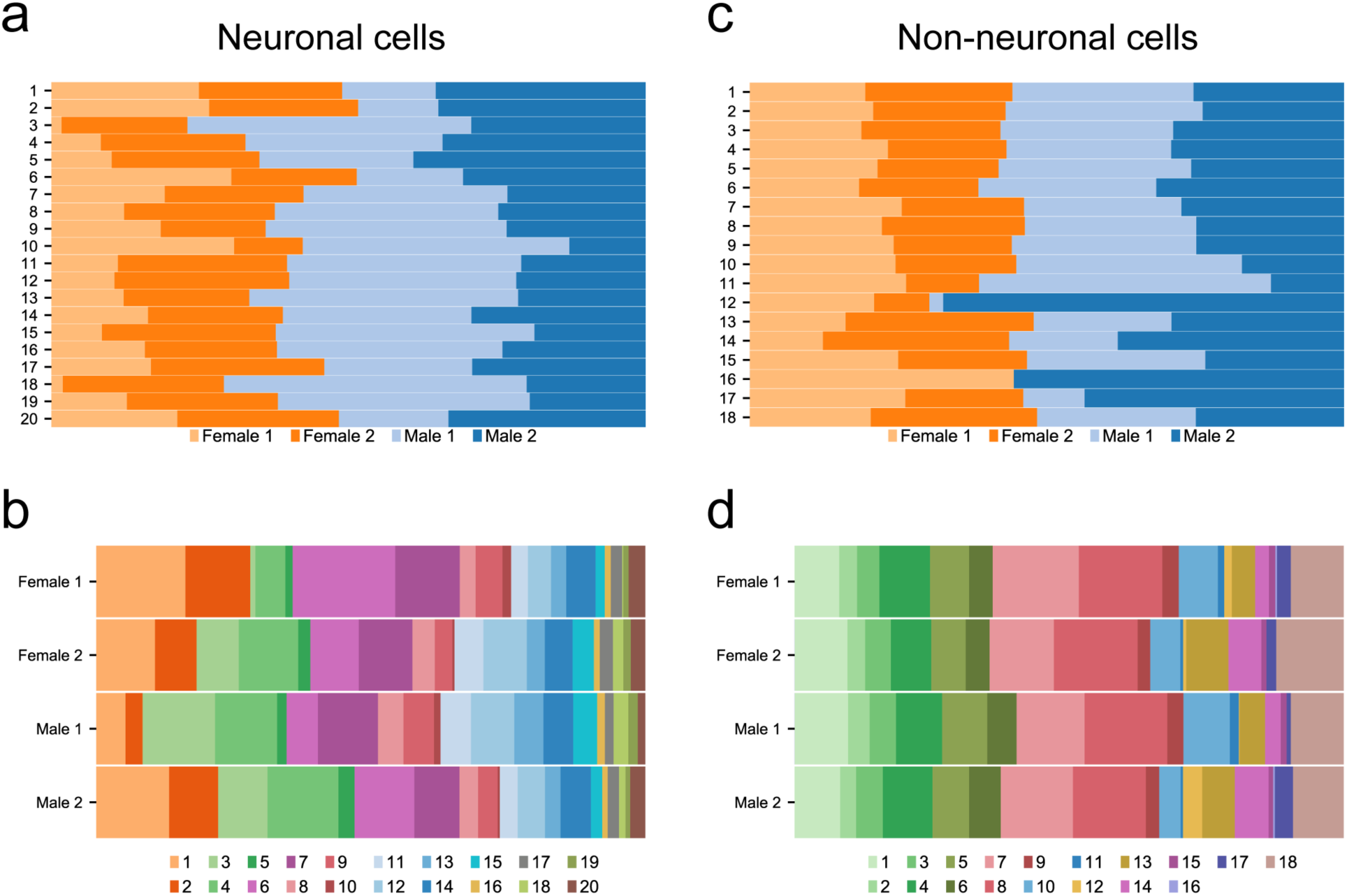
Proportion of cells derived from each sample contributing to each neuronal and non-neuronal cluster. **(a)** Proportion of cells (percent, %) from each sample contributing to each neuronal cluster; **(b)** from each neuronal cluster derived from each sample; **(c)** from each sample contributing to each non-neuronal cluster; and **(d)** from each non-neuronal cluster derived from each sample.

**Figure S3:**
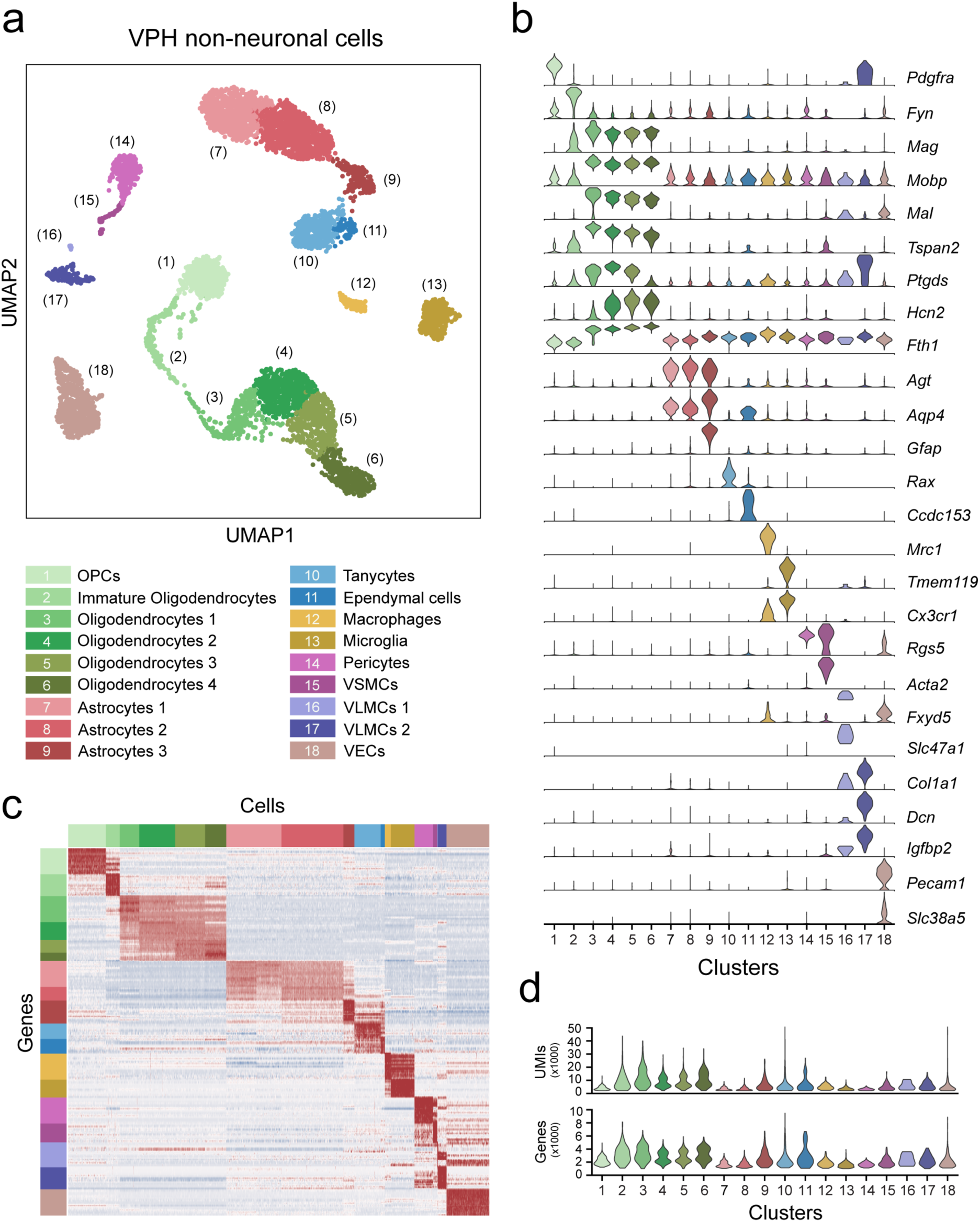
Classification of VPH non-neuronal populations. **(a)** UMAP plot showing unsupervised clustering of 18 VPH non-neuronal cells. **(b)** Violin plot showing the distribution of normalized expression of discriminatory marker genes in each cluster. Abbreviations: OPCs, oligodendrocyte precursor cells; VSMCs, vascular smooth muscle cells; VLMCs, vascular leptomeningeal cells; and VECs, vascular endothelial cells. **(c)** Heatmap showing scaled expression of discriminatory genes across all 18 non-neuronal clusters. **(d)** Violin plots showing the distribution in number of unique transcripts (upper) and number of genes in each non-neuronal cluster.

**Figure S4:**
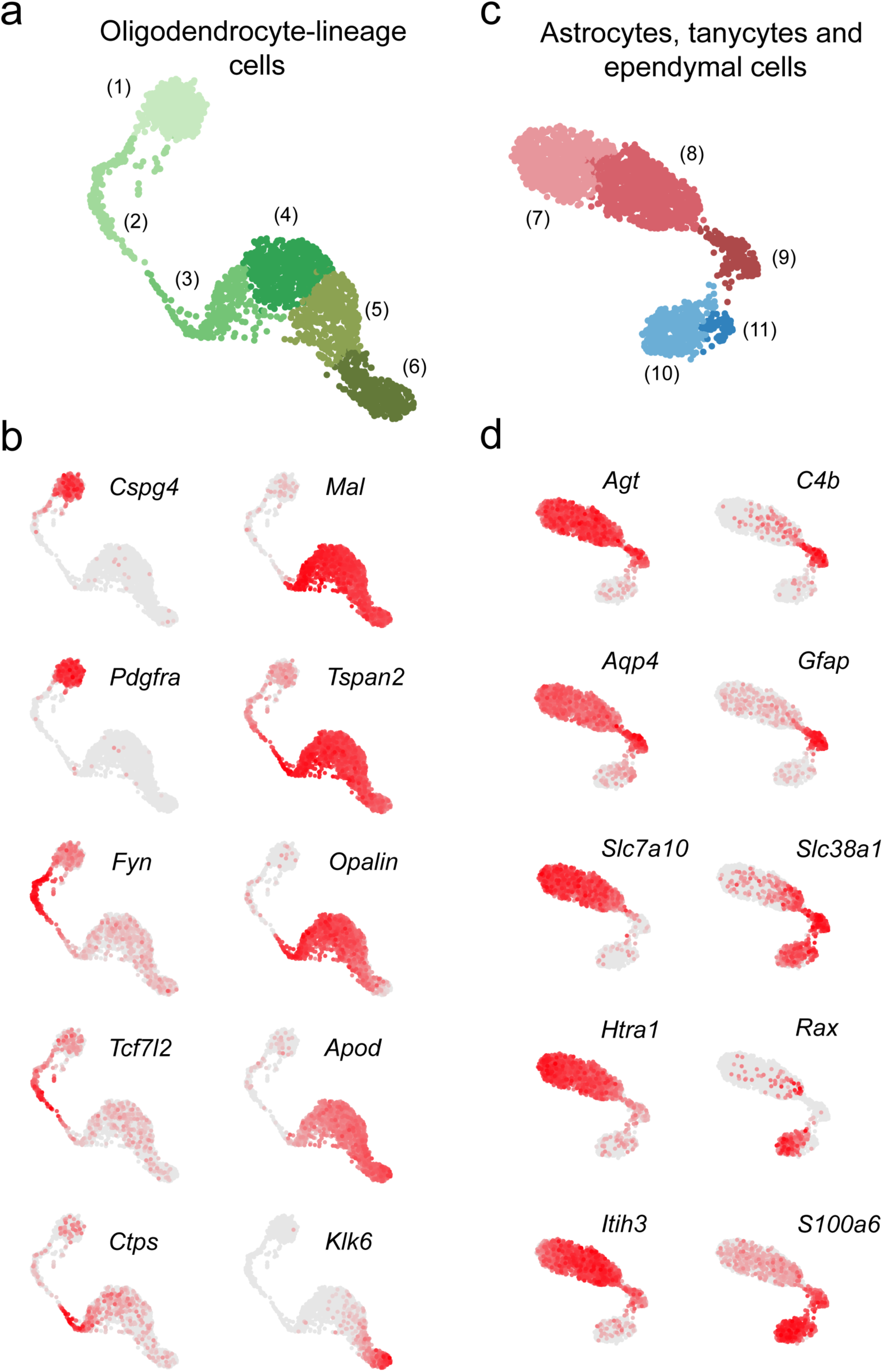
Key discriminatory marker genes of oligodendrocyte-lineage cells, astrocytes, tanycytes and ependymal cells. **(a)** UMAP plot showing just non-neuronal clusters 1-6 (detail of Fig. S3a). **(b)** Series of UMAP plots showing normalized expression of ten key discriminatory marker genes for subpopulations of oligodendrocyte lineage cells. **(c)** UMAP plot showing just non-neuronal clusters 7-11 (detail of Fig. S3a). **(d)** Series of UMAP plots showing normalized expression of ten key discriminatory marker genes for astrocyte subpopulations, tanycytes and ependymal cells.

**Figure S5:**
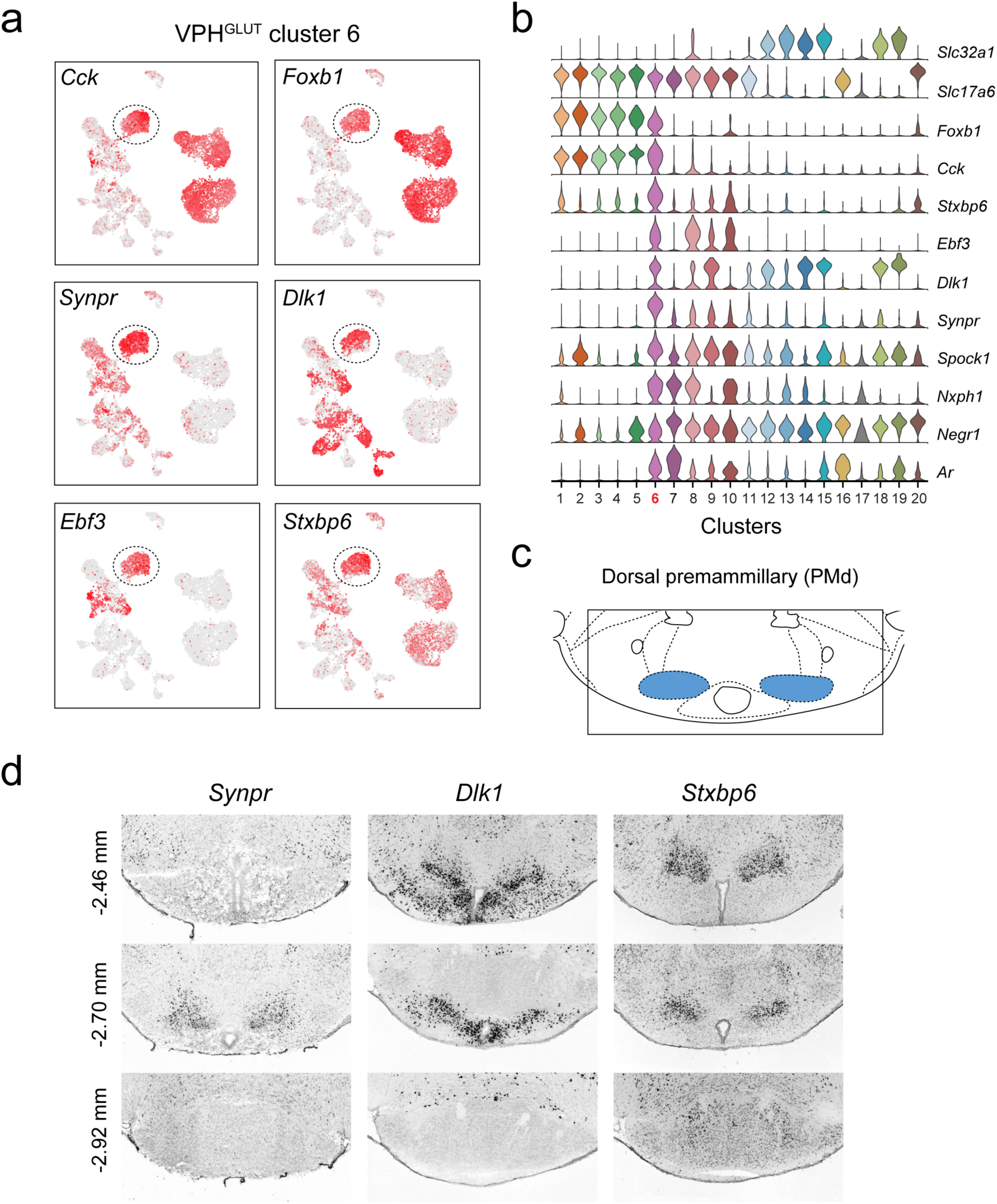
Identification of a population of putative *Synpr*+ PMd neurons. **(a)** UMAP plots showing normalized expression of *Cck, Foxb1* (shared with VPH^GLUT^ clusters 1-5), *Synpr, Dlk1, Ebf3* and *Stxbp6* (enriched in VPH^GLUT^ cluster 6). **(b)** Violin plot showing discriminatory marker genes enriched in VPH^GLUT^ cluster 6 following *Slc32a1* and *Slc17a6* (top). **(c)** Mouse brain atlas schematic, modified from Paxinos and Franklin, 2012, showing the PMd in a coronal section at distance from bregma of −2.70 mm (top). **(d)** ISH images for three anterior to posterior coronal sections (approximate distance from bregma −2.46, −2.80 and −2.92 mm) for *Synpr* (left), *Dlk1* (middle) and *Stxbp6* (right) from the ABA ^22^ (bottom). In each case, expression appears to be enriched in the PMd in anterior sections and largely absent in the MB in the posterior section.

**Figure S6:**
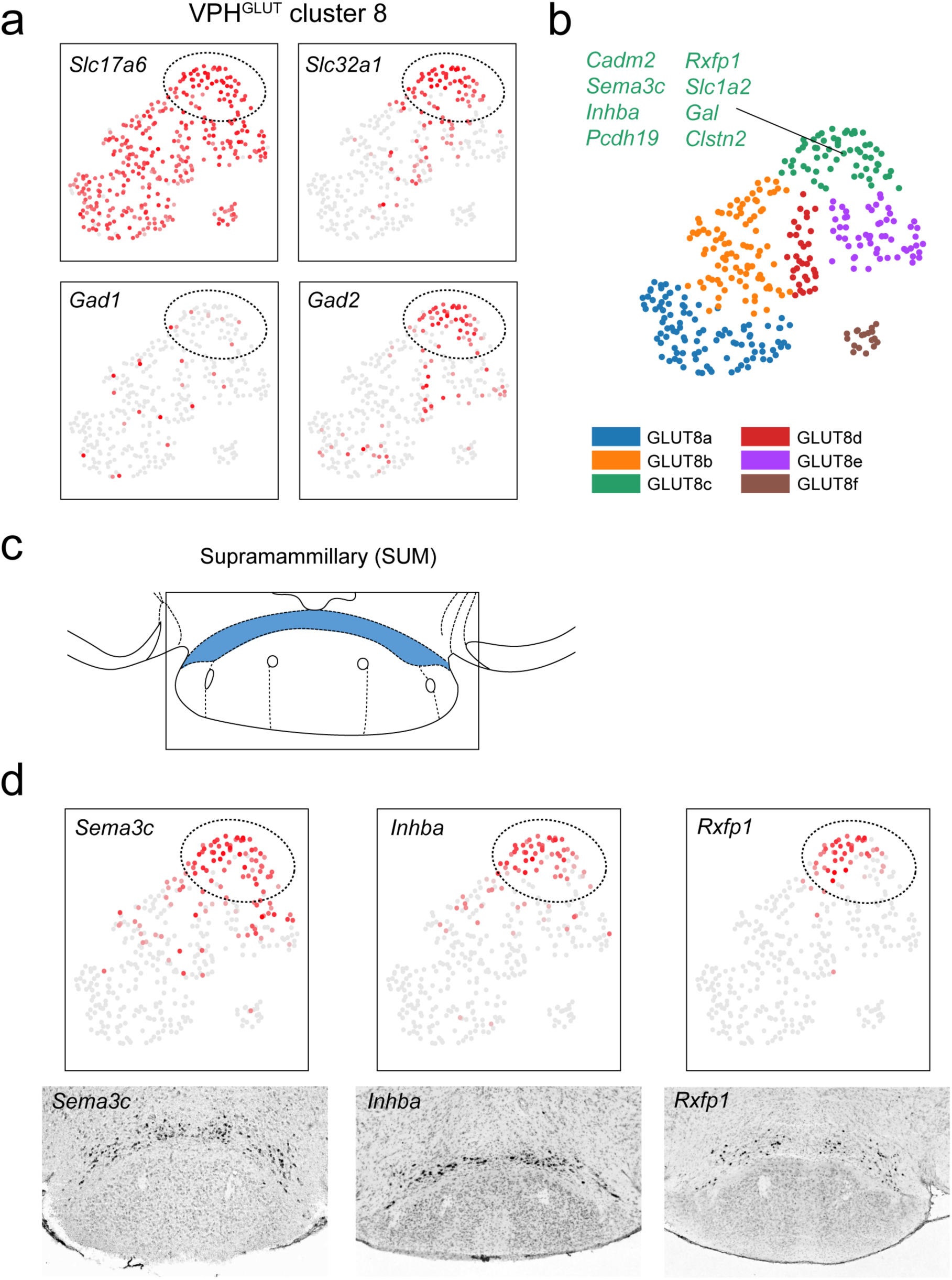
Identification of a population of putative dual phenotype GABA/glutamate *Sema3c*+ neurons in the SUM. **(a)** UMAP plots of only VPH^GLUT^ cluster 8, showing normalized expression of *Slc17a6, Slc32a1, Gad1* and *Gad2*. A region of cluster 8 (circled) was enriched in cells that were *Slc17a6*+, *Slc32a1*+, *Gad2*+ but *Gad1*-. **(b)** Further clustering of VPH^GLUT^ cluster 8 revealed six subclusters. Subcluster GLUT8c (green) expressed a suite of discriminatory genes including *Sema3c, Inhba* and *Rxfp1*. **(c)** Mouse brain atlas schematic, modified from Paxinos and Franklin, 2012, showing the SUM (or retromammillary nucleus/RM) in a coronal section at distance from bregma of −3.08 mm. **(d)** UMAP plots of only VPH^GLUT^ cluster 8, showing normalized expression of *Sema3c, Inhba* and *Rxfp1* (top). ISH images for *Sema3c, Inhba* and *Rxfp1* from the ABA ^22^ (bottom) showing localization to the posterior SUM.

**Figure S7:**
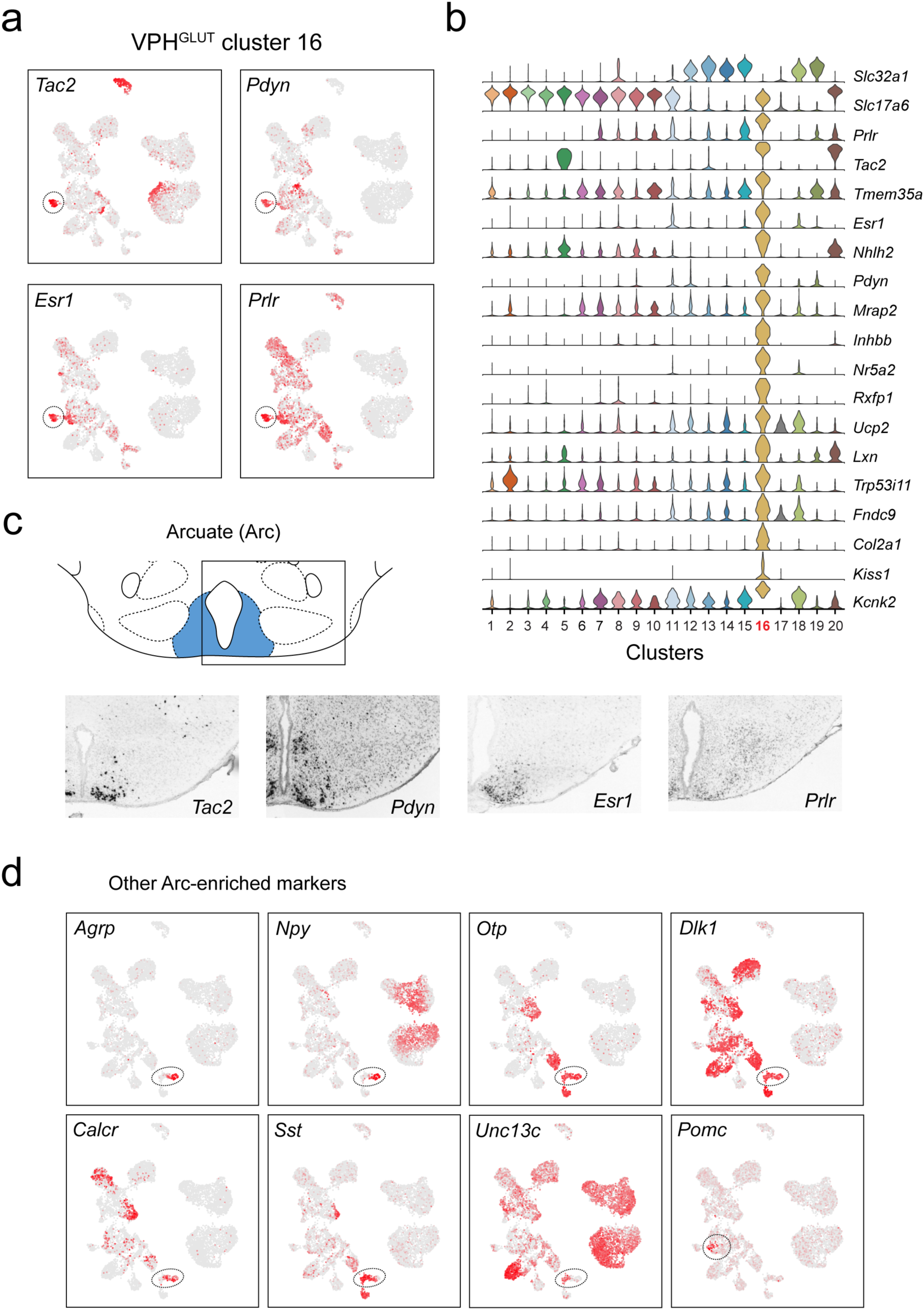
Identification of a population of putative Arc *Tac2*+ KNDy neurons and other Arc-enriched marker genes. **(a)** UMAP plots showing normalized expression of *Tac2, Pdyn, Esr1* and *Prlr* in VPH^GLUT^ cluster 16 (circled). **(b)** Violin plot showing discriminatory marker genes enriched in VPH^GLUT^ cluster 16 following *Slc32a1* and *Slc17a6* (top). **(c)** Mouse brain atlas schematic, modified from Paxinos and Franklin, 2012, showing the Arc (both lateroposterior and medial posterior subregions) in a coronal section at distance from bregma of −2.46 mm (top). ISH images for *Tac2, Pdyn, Esr1* and *Prlr* from the ABA ^22^ (bottom). **(d)** UMAP plots showing normalized expression of eight selected Arc-enriched markers in either VPH^GABA^ cluster 18 (circled in *Agrp, Npy, Otp, Dlk1, Calcr, Sst* and *Unc13c*) or VPH^GLUT^ cluster 11 (circled in *Pomc*).

**Figure S8:**
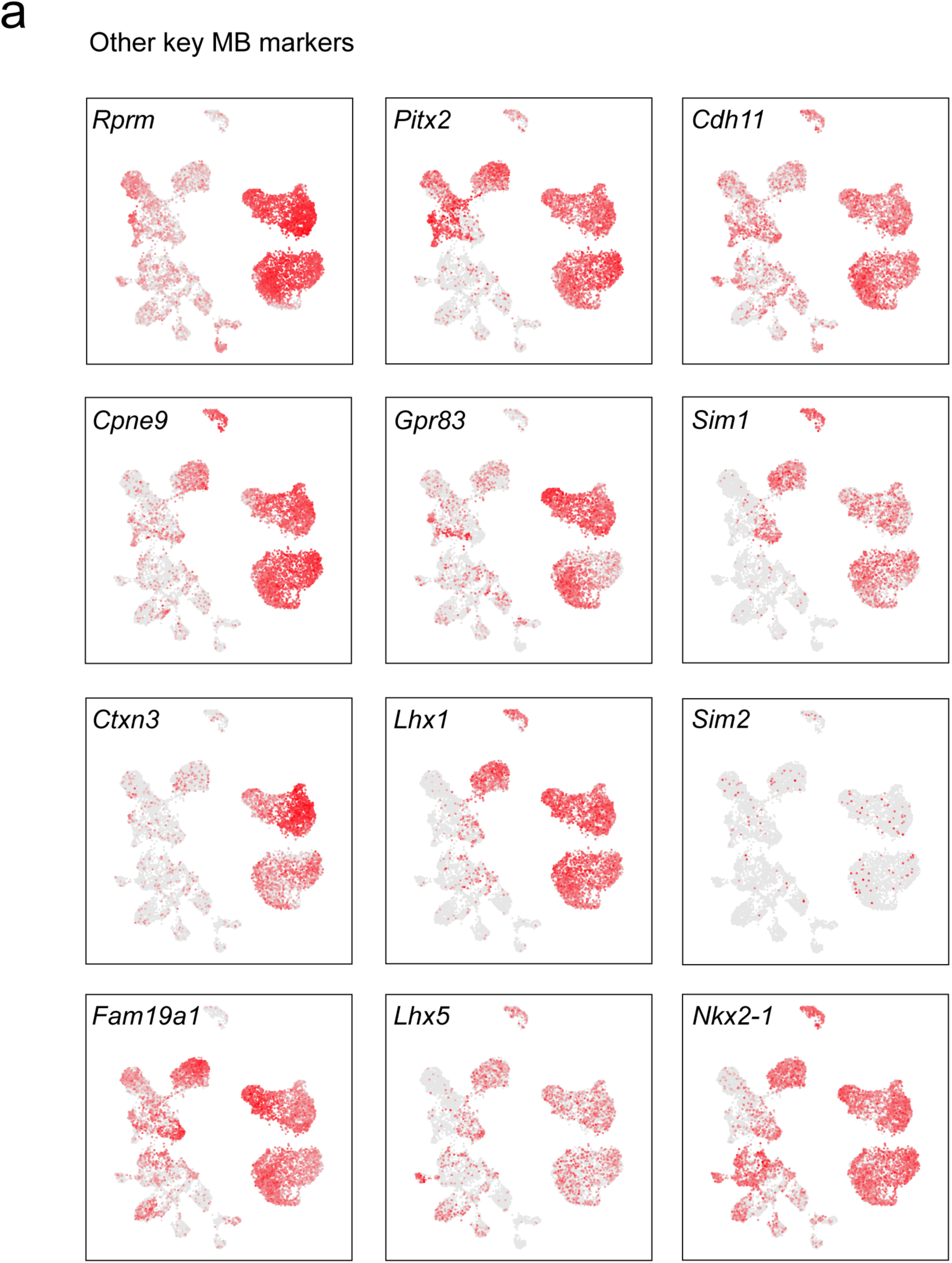
Other key discriminatory markers enriched in the MB. **(a)** UMAP plots showing normalized expression of discriminatory markers either identified in the current analysis or known to have a role in the development of the MB (VPH^GLUT^ cluster 1-5 and 20) and PMd (VPH^GLUT^ cluster 6).

## Notes

### Competing Interest Statement

The authors have declared no competing interest.

## REFERENCES

1. Saper, C. B. & Lowell, B. B. The hypothalamus. Curr. Biol. 24, R1111–R1116 (2014).

2. Simerly, R. B. Organization of the Hypothalamus. in The Rat Nervous System: Fourth Edition (2015). doi:10.1016/B978-0-12-374245-2.00013-9

3. Card, J. P. & Swanson, L. W. The Hypothalamus: An Overview of Regulatory Systems. in Fundamental Neuroscience: Fourth Edition (2013). doi:10.1016/B978-0-12-385870-2.00033-0

4. Papez, J. W. A proposed mechanism of emotion. Arch. Neurol. Psychiatry (1937). doi:10.1001/archneurpsyc.1937.02260220069003

5. Vann, S. D. & Aggleton, J. P. The mammillary bodies: Two memory systems in one? Nat. Rev. Neurosci. 5, 35–44 (2004).

6. Vann, S. D. Re-evaluating the role of the mammillary bodies in memory. Neuropsychologia 48, 2316–2327 (2010).

7. Aggleton, J. P. et al. Hippocampal-anterior thalamic pathways for memory: Uncovering a network of direct and indirect actions. Eur. J. Neurosci. 31, 2292–2307 (2010).

8. Dillingham, C. M., Frizzati, A., Nelson, A. J. D. & Vann, S. D. How do mammillary body inputs contribute to anterior thalamic function? Neurosci. Biobehav. Rev. 54, 108–119 (2015).

9. Vann, S. D. & Nelson, A. J. D. *The mammillary bodies and memory: More than a hippocampal relay*. Progress in Brain Research 219, (Elsevier B.V., 2015).

10. Canteras, N. S., Kroon, J. A. V., Do-Monte, F. H. M., Pavesi, E. & Carobrez, A. P. Sensing danger through the olfactory system: The role of the hypothalamic dorsal premammillary nucleus. Neurosci. Biobehav. Rev. 32, 1228–1235 (2008).

11. Donato, J. & Elias, C. F. C. F. The ventral premammillary nucleus links metabolic cues and reproduction. Front. Endocrinol. (Lausanne). 2, 1–10 (2011).

12. Leshan, R. L. & Pfaff, D. W. The hypothalamic ventral premammillary nucleus: A key site in leptin’s regulation of reproduction. J. Chem. Neuroanat. 61, 239–247 (2014).

13. Luppi, P. H., Billwiller, F. & Fort, P. Selective activation of a few limbic structures during paradoxical (REM) sleep by the claustrum and the supramammillary nucleus: evidence and function. Curr. Opin. Neurobiol. 44, 59–64 (2017).

14. Pan, W. X. & McNaughton, N. The supramammillary area: Its organization, functions and relationship to the hippocampus. Prog. Neurobiol. 74, 127–166 (2004).

15. Vertes, R. P. *Major diencephalic inputs to the hippocampus: Supramammillary nucleus and nucleus reuniens. Circuitry and function*. Progress in Brain Research 219, (Elsevier B.V., 2015).

16. Graebner, A. K., Iyer, M. & Carter, M. E. Understanding how discrete populations of hypothalamic neurons orchestrate complicated behavioral states. Front. Syst. Neurosci. 9, 111 (2015).

17. Andermann, M. L. & Lowell, B. B. Toward a Wiring Diagram Understanding of Appetite Control. Neuron 95, 757–778 (2017).

18. Lehman, M. N., Coolen, L. M. & Goodman, R. L. Minireview: Kisspeptin/neurokinin B/dynorphin (KNDy) cells of the arcuate nucleus: A central node in the control of gonadotropin-releasing hormone secretion. Endocrinology 151, 3479–3489 (2010).

19. Brown, R. E., Stevens, D. R. & Haas, H. L. The physiology of brain histamine. 63, 637–672 (2001).

20. Haas, H. L., Sergeeva, O. A. & Selbach, O. Histamine in the nervous system. Physiol. Rev. 88, 1183–241 (2008).

21. Panula, P. & Nuutinen, S. The histaminergic network in the brain: basic organization and role in disease. Nat. Rev. Neurosci. 14, 472–487 (2013).

22. Lein, E. S. et al. Genome-wide atlas of gene expression in the adult mouse brain. Nature 445, 168–176 (2007).

23. Zheng, G. X. Y. et al. Massively parallel digital transcriptional profiling of single cells. Nat. Commun. 8, (2017).

24. Paxinos, G. et al. Paxinos and Franklin’s the Mouse Brain in Stereotaxic Coordinates. Amsterdam, Academic Press (2012).

25. Marques, S. et al. Oligodendrocyte heterogeneity in the mouse juvenile and adult central nervous system. Science (80-.). 352, 1326–1329 (2016).

26. Zeisel, A. et al. Molecular Architecture of the Mouse Nervous System. Cell 174, 999-1014.e22 (2018).

27. Saunders, A. et al. Molecular Diversity and Specializations among the Cells of the Adult Mouse Brain. Cell 174, 1015-1030.e16 (2018).

28. Shimada, S. et al. Coexistence of substance P and neurotensin-like peptides in single neurons of the rat hypothalamus. Peptides 9, 71–76 (1988).

29. Larsen, P. J. Distribution of substance P-immunoreactive elements in the preoptic area and the hypothalamus of the rat. J. Comp. Neurol. 316, 287–313 (1992).

30. Vincent, S. R. & Kimura, H. Histochemical mapping of nitric oxide synthase in the rat brain. Neuroscience 46, 755–784 (1992).

31. Ross, R. A. et al. PACAP neurons in the ventral premammillary nucleus regulate reproductive function in the female mouse. Elife 7, 1–18 (2018).

32. Zoli, M., Agnati, L. F., Tinner, B., Steinbusch, H. W. M. & Fuxe, K. Distribution of dopamine-immunoreactive neurons and their relationships to transmitter and hypothalamic hormone-immunoreactive neuronal systems in the rat mediobasal hypothalamus. A morphometric and microdensitometric analysis. J. Chem. Neuroanat. 6, 293–310 (1993).

33. Meister, B. & Elde, R. Dopamine transporter mRNA in neurons of the rat hypothalamus. Neuroendocrinology 58, 388–395 (1993).

34. Soden, M. E. et al. Genetic Isolation of Hypothalamic Neurons that Regulate Context-Specific Male Social Behavior. Cell Rep. 16, 304–313 (2016).

35. Stagkourakis, S. et al. A neural network for intermale aggression to establish social hierarchy. Nat. Neurosci. 21, 834–842 (2018).

36. Boulland, J. L. et al. Vesicular glutamate and GABA transporters sort to distinct sets of vesicles in a population of presynaptic terminals. Cereb. Cortex 19, 241–248 (2009).

37. Soussi, R., Zhang, N., Tahtakran, S., Houser, C. R. & Esclapez, M. Heterogeneity of the supramammillary-hippocampal pathways: Evidence for a unique GABAergic neurotransmitter phenotype and regional differences. Eur. J. Neurosci. 32, 771–785 (2010).

38. Pedersen, N. P. et al. Supramammillary glutamate neurons are a key node of the arousal system. Nat. Commun. 8, (2017).

39. Hashimotodani, Y., Karube, F., Yanagawa, Y., Fujiyama, F. & Kano, M. Supramammillary Nucleus Afferents to the Dentate Gyrus Co-release Glutamate and GABA and Potentiate Granule Cell Output. Cell Rep. 25, 2704-2715.e4 (2018).

40. Moore, A. M., Coolen, L. M., Porter, D. T., Goodman, R. L. & Lehman, M. N. KNDy cells revisited. Endocrinology 159, 3219–3234 (2018).

41. Harter, C. J. L., Kavanagh, G. S. & Smith, J. T. The role of kisspeptin neurons in reproduction and metabolism. J. Endocrinol. 238, R173–R183 (2018).

42. Chen, R., Wu, X., Jiang, L. & Zhang, Y. Single-Cell RNA-Seq Reveals Hypothalamic Cell Diversity. Cell Rep. 18, 3227–3241 (2017).

43. Campbell, J. N. et al. A molecular census of arcuate hypothalamus and median eminence cell types. Nat. Neurosci. 20, 484–496 (2017).

44. Mickelsen, L. E. et al. Single-cell transcriptomic analysis of the lateral hypothalamic area reveals molecularly distinct populations of inhibitory and excitatory neurons. Nat. Neurosci. 22, 642–656 (2019).

45. Morales-Delgado, N. et al. Topography of somatostatin gene expression relative to molecular progenitor domains during ontogeny of the mouse hypothalamus. Front. Neuroanat. 5, 1–15 (2011).

46. Luo, S. X. et al. Regulation of feeding by somatostatin neurons in the tuberal nucleus. Science (80-.). (2018). doi:10.1126/science.aar4983

47. Ericson, H., Watanabe, T. & Köhler, C. Morphological analysis of the tuberomammmillary nucleus in the rat brain: Delineation of subgroups with antibody again L-histidine decarboxylase as a marker. J. Comp. Neurol. 263, 1–24 (1987).

48. Inagaki, N. et al. Organization of histaminergic fibers in the rat brain. J. Comp. Neurol. 273, 283–300 (1988).

49. Eriksson, K. S. et al. The type III neurofilament peripherin is expressed in the tuberomammillary neurons of the mouse. BMC Neurosci. 9, 26 (2008).

50. Vincent, S. R., Hökfelt, T., Skirboll, L. R. & Wu, J. Y. Hypothalamic γ-aminobutyric acid neurons project to the neocortex. Science (80-.). (1983). doi:10.1126/science.6857253

51. Takeda, N. et al. Immunohistochemical evidence for the coexistence of histidine decarboxylase-like and glutamate decarboxylase-like immunoreactivities in nerve cells of the magnocellular nucleus of the posterior hypothalamus of rats. Proc. Natl. Acad. Sci. U. S. A. 81, 7647–7650 (1984).

52. Yu, X. et al. Wakefulness Is Governed by GABA and Histamine Cotransmission. Neuron 87, 164–178 (2015).

53. Venner, A. et al. Reassessing the Role of Histaminergic Tuberomammillary Neurons in Arousal Control. J. Neurosci. 39, 8929–8939 (2019).

54. Dillingham, C. M. & Vann, S. D. Why Isn’t the Head Direction System Necessary for Direction? Lessons From the Lateral Mammillary Nuclei. Front. Neural Circuits 13, 1–10 (2019).

55. Kim, D. W. et al. The cellular and molecular landscape of hypothalamic patterning and differentiation. bioRxiv 657148 (2020). doi:10.1101/657148

56. Kimura, S. et al. The T/ebp null mouse: Thyroid-specific enhancer-binding protein is essential for the organogenesis of the thyroid, lung, ventral forebrain, and pituitary. Genes Dev. 10, 60–69 (1996).

57. Bedont, J. L., Newman, E. A. & Blackshaw, S. Patterning, specification, and differentiation in the developing hypothalamus. Wiley Interdiscip. Rev. Dev. Biol. 4, 445–468 (2015).

58. Puelles, L. et al. Pallial and subpallial derivatives in the embryonic chick and mouse telencephalon, traced by the expression of the genes Dlx-2, Emx-1, Nkx-2.1, Pax-6, and Tbr-1. J. Comp. Neurol. 424, 409–438 (2000).

59. Martin, D. M., Skidmore, J. M., Fox, S. E., Gage, P. J. & Camper, S. A. Pitx2 distinguishes subtypes of terminally differentiated neurons in the developing mouse neuroepithelium. Dev. Biol. 252, 84–99 (2002).

60. Marion, J. F., Yang, C., Caqueret, A., Boucher, F. & Michaud, J. L. Sim1 and Sim2 are required for the correct targeting of mammillary body axons. Development 132, 5527–5537 (2005).

61. Shimogori, T. et al. A genomic atlas of mouse hypothalamic development. Nat. Neurosci. 13, 767–775 (2010).

62. Skidmore, J. M., Waite, M. R., Alvarez-Bolado, G., Puelles, L. & Martin, D. M. A novel TaulacZ allele reveals a requirement for Pitx2 in formation of the mammillothalamic tract. Genesis 50, 67–73 (2012).

63. Miquelajáuregui, A. et al. LIM homeobox protein 5 (Lhx5) is essential for mamillary body development. Front. Neuroanat. 9, 1–10 (2015).

64. Szabó, N. E., Haddad-Tóvolli, R., Zhou, X. & Alvarez-Bolado, G. Cadherins mediate sequential roles through a hierarchy of mechanisms in the developing mammillary body. Front. Neuroanat. 9, 1–17 (2015).

65. Ferran, J. L., Puelles, L. & Rubenstein, J. L. R. Molecular codes defining rostrocaudal domains in the embryonic mouse hypothalamus. Front. Neuroanat. 9, 1–14 (2015).

66. Shibata, H. Topographic organization of subcortical projections to the anterior thalamic nuclei in the rat. J. Comp. Neurol. 323, 117–127 (1992).

67. Jankowski, M. M. et al. The anterior thalamus provides a subcortical circuit supporting memory and spatial navigation. Front. Syst. Neurosci. 7, 1–12 (2013).

68. Bubb, E. J., Kinnavane, L. & Aggleton, J. P. Hippocampal–diencephalic–cingulate networks for memory and emotion: An anatomical guide. Brain Neurosci. Adv. 1, 239821281772344 (2017).

69. Zakowski, W., Równiak, M. & Robak, A. Colocalization pattern of calbindin and cocaine- and amphetamine-regulated transcript in the mammillary body-anterior thalamic nuclei axis of the guinea pig. Neuroscience 260, 98–105 (2014).

70. Romanov, R. A. et al. Molecular interrogation of hypothalamic organization reveals distinct dopamine neuronal subtypes. Nat. Neurosci. 20, (2016).

71. Moffitt, J. R. et al. Molecular, spatial, and functional single-cell profiling of the hypothalamic preoptic region. Science (80-.). 362, (2018).

72. Rossi, M. A. et al. Obesity remodels activity and transcriptional state of a lateral hypothalamic brake on feeding. Science (80-.). 364, 1271–1274 (2019).

73. Kim, D. W. et al. Multimodal Analysis of Cell Types in a Hypothalamic Node Controlling Social Behavior. Cell 179, 713-728.e17 (2019).

74. van den Pol, A. N. Neuropeptide Transmission in Brain Circuits. Neuron 76, 98–115 (2012).

75. Vann, S. D. & Aggleton, J. P. Evidence of a spatial encoding deficit in rats with lesions of the mammillary bodies or mammillothalamic tract. J. Neurosci. 23, 3506–3514 (2003).

76. Vann, S. D. Dismantling the papez circuit for memory in rats. Elife 2013, 1–21 (2013).

77. Tsivilis, D. et al. A disproportionate role for the fornix and mammillary bodies in recall versus recognition memory. Nat. Neurosci. 11, 834–842 (2008).

78. Kopelman, M. D. The Korsakoff syndrome. British Journal of Psychiatry (1995). doi:10.1192/bjp.166.2.154

79. Kril, J. J. & Harper, C. G. Neuroanatomy and neuropathology associated with Korsakoff’s syndrome. Neuropsychol. Rev. 22, 72–80 (2012).

80. Kaestner, K. H., Schütz, G. & Monaghan, A. P. Expression of the winged helix genes fkh-4 and fkh-5 defines domains in the central nervous system 1. Mech. Dev. 55, 221–230 (1996).

81. Wehr, R., Mansouri, A., De Maeyer, T. & Gruss, P. Fkh5-deficient mice show dysgenesis in the caudal midbrain and hypothalamic mammillary body. Development 124, 4447–4456 (1997).

82. Radyushkin, K. et al. Genetic ablation of the mammillary bodies in the Foxb1 mutant mouse leads to selective deficit of spatial working memory. Eur. J. Neurosci. 21, 219–229 (2005).

83. Alvarez-Bolado, G., Zhou, X., Voss, A. K., Thomas, T. & Gruss, P. Winged helix transcription factor Foxb1 is essential for access of mammillothalamic axons to the thalamus. Development 127, 1029–1038 (2000).

84. Gomes, I. et al. Identification of GPR83 as the receptor for the neuroendocrine peptide PEN. Sci. Signal. 9, 1–15 (2016).

85. Pesini, P., Detheux, M., Parmentier, M. & Hökfelt, T. Distribution of a glucocorticoid-induced orphan receptor (JP05) mRNA in the central nervous system of the mouse. Mol. Brain Res. 57, 281–300 (1998).

86. E. Vollmer, L. et al. Attenuated stress-evoked anxiety, increased sucrose preference and delayed spatial learning in glucocorticoid-induced receptor-deficient mice. Genes, Brain Behav. (2013). doi:10.1111/j.1601-183X.2012.00867.x

87. Canteras, N. S. & Swanson, L. W. The dorsal premammillary nucleus: An unusual component of the mammillary body. Proc. Natl. Acad. Sci. U. S. A. 89, 10089–10093 (1992).

88. Cezario, A. F., Ribeiro-Barbosa, E. R., Baldo, M. V. C. & Canteras, N. S. Hypothalamic sites responding to predator threats - The role of the dorsal premammillary nucleus in unconditioned and conditioned antipredatory defensive behavior. Eur. J. Neurosci. 28, 1003–1015 (2008).

89. Blanchard, D. C. et al. Dorsal premammillary nucleus differentially modulates defensive behaviors induced by different threat stimuli in rats. Neurosci. Lett. 345, 145–148 (2003).

90. Allen, G. V. & Hopkins, D. A. Mamillary body in the rat: A cytoarchitectonic, golgi, and ultrastructural study. J. Comp. Neurol. 275, 39–64 (1988).

91. Allen, G. V. & Hopkins, D. A. Mamillary body in the rat: Topography and synaptology of projections from the subicular complex, prefrontal cortex, and midbrain tegmentum. J. Comp. Neurol. 286, 311–336 (1989).

92. Christiansen, K. et al. Complementary subicular pathways to the anterior thalamic nuclei and mammillary bodies in the rat and macaque monkey brain. Eur. J. Neurosci. 43, 1044–1061 (2016).

93. Bienkowski, M. S. et al. Integration of gene expression and brain-wide connectivity reveals the multiscale organization of mouse hippocampal networks. Nat. Neurosci. 21, 1628–1643 (2018).

94. Cembrowski, M. S. et al. Dissociable Structural and Functional Hippocampal Outputs via Distinct Subiculum Cell Classes. Cell 173, 1280-1292.e18 (2018).

95. Harris, J. A. et al. Hierarchical organization of cortical and thalamic connectivity. Nature 575, 195–202 (2019).

96. Phillips, J. W. et al. A repeated molecular architecture across thalamic pathways. Nat. Neurosci. 22, 1925–1935 (2019).

97. Callen, D. J. A., Black, S. E., Gao, F., Caldwell, C. B. & Szalai, J. P. Beyond the hippocampus: MRI volumetry confirms widespread limbic atrophy in AD. Neurology (2001). doi:10.1212/WNL.57.9.1669

98. Copenhaver, B. R. et al. The fornix and mammillary bodies in older adults with Alzheimer’s disease, mild cognitive impairment, and cognitive complaints: A volumetric MRI study. Psychiatry Res. - Neuroimaging 147, 93–103 (2006).

99. Grossi, D., Lopez, O. L. & Martinez, A. J. Mamillary bodies in Alzheimer’s disease. Acta Neurol. Scand. (1989). doi:10.1111/j.1600-0404.1989.tb03840.x

100. Baloyannis, S. J., Mavroudis, I., Baloyannis, I. S. & Costa, V. G. Mammillary Bodies in Alzheimer’s Disease: A Golgi and Electron Microscope Study. Am. J. Alzheimers. Dis. Other Demen. 31, 247–256 (2016).

101. Canter, R.G. et al. 3D mapping reveals network-specific amyloid progression and subcortical susceptibility in mice. Commun. Biol. 2, 1–12 (2019).

102. Oakley, H. et al. Intraneuronal β-amyloid aggregates, neurodegeneration, and neuron loss in transgenic mice with five familial Alzheimer’s disease mutations: Potential factors in amyloid plaque formation. J. Neurosci. (2006). doi:10.1523/JNEUROSCI.1202-06.2006

103. Saper, C. B. & German, D. C. Hypothalamic pathology in Alzheimer’s disease. Neurosci. Lett. 74, 364–370 (1987).

104. Alraksinen, M. S. et al. Histamine neurons in human hypothalamus: Anatomy in normal and alzheimer diseased brains. Neuroscience 44, 465–481 (1991).

105. Nakamura, S. et al. Loss of large neurons and occurrence of neurofibrillary tangles in the tuberomammillary nucleus of patients with Alzheimer’s disease. Neurosci. Lett. 151, 196–199 (1993).

106. Oh, J. et al. Profound degeneration of wake-promoting neurons in Alzheimer’s disease. Alzheimer’s Dement. 15, 1253–1263 (2019).

107. Spira, A. P., Chen-Edinboro, L. P., Wu, M. N. & Yaffe, K. Impact of sleep on the risk of cognitive decline and dementia. Curr. Opin. Psychiatry 27, 478–483 (2014).

108. Vong, L. et al. Leptin Action on GABAergic Neurons Prevents Obesity and Reduces Inhibitory Tone to POMC Neurons. Neuron 71, 142–154 (2011).

109. Leinninger, G. M. et al. Leptin action via neurotensin neurons controls orexin, the mesolimbic dopamine system and energy balance. Cell Metab. 14, 313–23 (2011).

110. Wolf, F. A., Angerer, P. & Theis, F. J. SCANPY: Large-scale single-cell gene expression data analysis. Genome Biol. (2018). doi:10.1186/s13059-017-1382-0

111. Macosko, E. Z. et al. Highly parallel genome-wide expression profiling of individual cells using nanoliter droplets. Cell 161, 1202–1214 (2015).

112. Korsunsky, I. et al. Fast, sensitive and accurate integration of single-cell data with Harmony. Nat. Methods 16, 1289–1296 (2019).

